# Assessing the role of inversions in maintaining genomic differentiation after secondary contact: local adaptation, genetic incompatibilities, and drift

**DOI:** 10.1101/2020.08.25.267369

**Authors:** Marina Rafajlović, Jordi Rambla, Jeffrey L. Feder, Arcadi Navarro, Rui Faria

## Abstract

Due to their effects on reducing recombination, chromosomal inversions may play an important role in speciation by establishing and/or maintaining linked blocks of genes causing reproductive isolation (RI) between populations. These views fit empirical data indicating that inversions typically harbour loci involved in RI. However, previous computer simulations of infinite populations with 2-4 loci involved in RI implied that, even with gene flux as low as 10^−8^ between alternative arrangements, inversions may not have large, qualitative advantages over collinear regions in maintaining population differentiation after secondary contact. Here, we report that finite population sizes can help counteract the homogenizing consequences of gene flux, especially when several fitness-related loci reside within the inversion. In these cases, the persistence time of differentiation after secondary contact can be similar to when gene flux is absent, and notably longer than the persistence time without inversions. Thus, despite gene flux, population differentiation may be maintained for up to 100,000 generations, during which time new incompatibilities and/or local adaptations might accumulate and facilitate progress towards speciation. How often these conditions are met in nature remains to be determined.

## Introduction

Chromosomal rearrangements, specifically inversions, have been suggested to play key roles in adaptation and speciation practically since the beginning of genetics (Sturtevant 1917; reviewed by Jackson *et al.* 2016). Inversions were first identified by their effects on suppressing recombination (Rieseberg 2001; Faria and Navarro 2010), and in the past 30 years, their role as recombination modifiers became central to many speciation models (Trickett and Butlin 1994; Rieseberg 2001; Noor et al. 2001).

In heterozygotes for one or more inversions (*i.e.*, heterokaryotypes), recombination can be severely reduced within the inverted regions (Sturtevant and Beadle 1936). Consequently, genetic variation within inversions is expected to resist homogenization by gene flow between populations fixed for alternative arrangements relative to collinear regions of the genome (Rieseberg 2001). Chromosomal inversions may reduce gene flow sufficiently to facilitate progress towards complete speciation (Trickett and Butlin 1994; Navarro and Barton 2003; Kirkpatrick and Barton 2006; Feder *et al.* 2011; Guerrero *et al.* 2012; Wellenreuther and Bernatchez 2018).

However, even when recombination due to single crossing over is reduced in heterokaryotypes within regions spanned by inversions, gene flux, the movement of DNA sequences between alternative karyotypes (Navarro *et al.* 1997a) is not completely eliminated. Gene flux can occur via gene conversion and/or double crossovers (Navarro *et al.* 1997a, 1997b; Stevison *et al.* 2011; Korunes and Noor 2016; Crown *et al.* 2018; Korunes and Noor 2019; Fuller *et al.* 2019; Faria *et al.* 2019b). Gene flux will tend to reduce genetic differentiation between chromosomes bearing different arrangements (Korunes and Noor 2016; Korunes and Noor 2019).

Homogenization will be faster when the genetic content within alternative arrangements is neutral than when different populations are fixed for alternative arrangements and the inversion contains loci that are under divergent selection or exhibit some form of genetic incompatibility (Guerrero *et al.* 2012; Fuller *et al.* 2019). In general, a reduction of gene flux would result in a reduction of gene flow between populations in the segments spanned by inversions.

In other words, inversion polymorphisms are semi-permeable barriers to gene flow and, thus, the question is: under which conditions do they play an effective role in adaptation, divergence, and speciation in the face of gene flow? Theoretical work about the evolutionary role of inversion polymorphisms has focused mainly on two-population models where either the alternative chromosomal arrangements themselves or their allelic content undergo divergent selection. Kirkpatrick and Barton (2006) and Charlesworth and Barton (2018) analyzed the conditions favoring the spread of an initially rare inversion, assuming that the inversion polymorphism captures two or more adaptive loci undergoing divergent selection. A general finding of these studies is that, by suppressing recombination, an inversion protects locally favorable combinations of alleles from mixing with maladapted alleles introduced by migration (except when migration is too strong, see Eq. A9 in Charlesworth and Barton, 2018). In addition, Nei et al. (1967) have shown that, under a mutation-selection equilibrium model, an inversion can also invade without requiring divergent selection, as long as it harbors a low number of non-deleterious alleles.

Although these studies assumed populations of infinite size, it is straightforward to deduce that, in populations of finite size diverging in the face of gene flow by the accumulation of new adaptive mutations, a spreading inversion that happens to capture two or more locally favorable alleles will be an effective mechanism protecting these alleles from stochastic loss (Rafajlović *et al.* 2016), thereby facilitating divergence. Furthermore, genetic patterns at neutral loci linked to an inversion polymorphism in pairs of divergent populations subject to migration have been assessed by Guerrero *et al.* (2012) under a coalescent framework. Their results suggest that inversion polymorphisms can store differentiation at linked neutral loci depending on the age of the inversion, on gene flux relative to the recombination rate in homokaryotypes, and on migration relative to selection.

While a significant amount of work has been devoted to understanding the evolutionary role of inversion polymorphisms when divergent selection acts between populations, less attention has been paid to alternative models that do not involve geographically divergent selection *per se* and yet allow for the establishment of partial barriers to gene flow. Examples include models involving neutral and universally beneficial alleles, with or without incompatibilities (Noor *et al.* 2001; Navarro and Barton 2003), or modified versions of such models (cf. Feder and Nosil 2009). Such models are relevant when considering pairs of populations without any obvious ecological trade-offs. Empirical data supporting models with locally beneficial and neutral alleles (so-called *conditional-neutrality models*) can be found in Anderson et al. (2012) (see also references therein). Models involving universally beneficial alleles with genetic incompatibilities have also been suggested in the literature, for instance in studies of *Drosophila pseudoobscura* and *D. persimilis* (Noor *et al.* 2001; Kulathinal *et al.* 2009). These two species exhibit higher divergence in genomic regions fixed for alternative arrangements, and precisely these regions harbor loci contributing to the reproductive isolation (RI) between these species (Noor *et al.* 2001; Kulathinal *et al.* 2009). These empirical findings may be explained (though not exclusively, see Fuller *et al.* 2018) by invoking genetic incompatibilities: whereas genetic incompatibilities established in each species could have been eliminated from collinear regions of the genome after the species’ secondary contact, incompatibilities might have been maintained within inversion polymorphisms due to suppressed recombination in heterokaryotypes (Noor *et al.* 2001; Ortiz-Barrientos *et al.* 2002; Navarro and Barton 2003). Thus, inversion polymorphisms could have played a crucial role in the persistence of these two species upon their secondary contact, allowing for additional reproductive barriers (*i.e.*, reinforcement) to accumulate thereafter (Noor *et al.* 2001).

To understand the interplay between gene flux and the type of selection acting on variation within an inversion polymorphism, Feder and Nosil (2009) implemented simulations of secondary contact between two strongly differentiated populations with initially fixed alternative arrangements. After testing five different models ‒some involving local adaptation, with and without trade-offs, and others involving genetic incompatibilities – the authors showed that, in the absence of divergent selection, there are conditions under which inversions may help retain species differences following secondary contact compared to collinear gene regions lacking rearrangements. However, the authors also showed that large, qualitative differences between inversions versus collinear regions will not persist for long time periods (tens of thousands of generations) even in the presence of low levels of gene flux between alternative arrangements (*e.g.*, 10^−8^), and when migration between populations is not too weak compared to selection at loci causing RI. Given that estimates of gene conversion have been recently shown to be potentially as high as 10^−5^ to 2.5·10^−5^ between rearrangements in *Drosophila* (Korunes and Noor 2019), this raises the question as to the efficacy of inversions in maintaining prolonged diffentiation following secondary contact in cases without divergent selection.

Feder and Nosil (2009) focused on inversions containing only two or four loci and on deterministic scenarios (with populations of infinite size). Here, we present a series of simulations that extend the framework of Feder and Nosil (2009) for the cases of locally adaptive alleles and genetic incompatibilities (Models 2 and 4 in their study). Our goal is to test how the effects of more than two loci and of finite population sizes may influence the erosion of differentiation within an inversion polymorphism after secondary contact, paying particular attention to the recent empirical results implying higher rates of gene flux between chromosomal rearrangements than previously thought (Korunes and Noor 2019).

We show that gene flux between alternative chromosomal rearrangements can be retarded by the finite size of populations (with up to 100,000 individuals in some cases) coming into secondary contact. This is due to a longer waiting time for actual gene flux events to be realized, and also, to some extent, due to the increased probability of the loss of favored allele combinations by random genetic drift. In particular, we find that under these conditions, inversions can retain longer-lived differentiation even with relatively high levels of gene flux (2 · 10^−4^ per gamete per generation) when many loci of small fitness effects reside within the inversion polymorphism. Importantly, such inversion polymorphisms can protect population differentiation for tens of thousands of generations (up to 100,000, or longer in some cases) after secondary contact. This time span may provide a window of opportunity for additional differences to accumulate (further local adaptation, more incompatibilities, reinforcement, etc.), facilitating further progress towards speciation. How often these conditions are met for hybridizing populations in nature remains to be determined. Nevertheless, our results do offer a region of parameter space where inversions not containing loci causing fitness trade-offs (divergent selection) in alternate habitats may play an enhanced role in facilitating speciation than collinear regions following secondary contact.

## Model and Methods

To assess the role of multiple loci and finite population sizes in maintaining population differentiation after secondary contact, we used individual-based computer simulations of two diploid populations with finite, constant population size (denoted by *N* below) and discrete, non-overlapping generations. At the start of each simulation, the two populations (referred to as Population 1 and Population 2) were fixed for different chromosomal arrangements. For convenience, we refer to the arrangement in Population 1 as *standard* and to the other one as *inverted*. We modelled *2L* bi-allelic loci within the region polymorphic for the inversion (see below) and assumed that at the start of each simulation the two populations were fixed for alternative alleles at each locus. At generation 0, both populations consisted of adult virgin individuals coming into secondary contact. Thereafter, adult virgin individuals underwent migration, followed by soft selection, recombination and mating (always locally in each population). We neglected mutations throughout. Migration from one population to the other occurred with a per-generation per-individual probability of *m*. After migration, we applied soft fecundity selection within each population, so that the number of gametes that an adult individual contributed to the pool of offspring was binomially distributed with the number of trials equal to *2N*, and the probability equal to the fitness of the individual in the population where it ended up after migration, relative to the total fitness of all adult individuals in this population.

In homokaryotypes (either for the standard or for the inverted chromosomal arrangement), consecutive loci in the simulated genomic region underwent recombination with a per-gamete per-generation probability of *r*_2*L*_. We modelled the recombination rate *r*_2*L*_ in two different ways, that we explain below.

First, we focused on inversions of the same size in terms of their total map distance. Throughout, we refer to this version of the model as the *conserved-size model*. Here, we chose the recombination rate *r*_2*L*_ in such a way that it was smaller when the number of fitness-related loci (*i.e., 2L*) in the inversion polymorphism was larger, and that the total recombination distance (denoted by *r* below) spanned by the inversion was independent of the number of loci it contained. In particular, and in line with Feder and Nosil (2009), when 2*L* = 2, we set *r*_2_ = *r* = 0.1, whereas for 2*L* > 2, we used *r*_2*L*_ = 1 − (1 − *r*)^1/(2*L*-1)^. In addition, due to our focus on the inversion, gene flux between the alternative arrangements in heterokaryotypes occurred with a per-gamete per-generation probability *r*_Inv_, that was independent of the number of loci 2*L*. Gene flux was implemented essentially as in Feder and Nosil (2009), except that we assumed that gene flux occurred 50% of the time by gene conversion and 50% by double crossover (whereas the corresponding ratio in Feder and Nosil (2009) was 30:70). This difference takes into account a recent study by Korunes and Noor (2019) showing that double crossover in hybrids between *Drosophila pseudoobscura* and *D. persimilis* result in slightly lower gene flux per site and generation than gene conversion; but the reverse is observed for double crossovers spanning a region larger than 1 Mb. Since in our simulations the ratio between the number of exchanged alleles due to double crossover and gene conversion varies depending on how many loci are exchanged in each double crossover (see below), we assumed equal rates of double crossover and gene conversion to avoid any bias towards one or the other process. In our model, a double crossover was assumed to occur within a region containing any number between 1 and *2L* of consecutive loci, each combination being equally likely. Gene conversion events were modelled so that an allele at a given locus at a given chromosome was altered to the allele at the same locus residing at the homologous chromosome. If the two alleles were identical prior to gene conversion, they remained identical also afterwards. Gene conversion events occurred at a single locus, each locus being equally likely.

In a second version of the model, we assumed that the size of the inversion increased with the number of fitness-related loci within it. We refer to this version of the model as the *increasing-size model*. Here, we set the recombination rate *r*_2*L*_ between each pair of consecutive loci in homokaryotypes to be independent of the number of fitness-related loci within the inversion, *i.e.*, *r*_2*L*_ = *r* = 0.1. In this version of the model, the total rate of gene flux in heterokaryotypes (denoted by *r*_Inv,2*L*_ below) was assumed to depend linearly on the number of fitness-related loci, *i.e.*, *r*_Inv,2*L*_ = *r*_Inv_*L*, where *r*_Inv_ had the same value as in the *conserved-size* model explained above. Note that this means that when the inversion contained exactly two fitness-related loci (*i.e.*, 2*L* = 2), *the conserved-size* model and the *increasing-size* model were equivalent, whereas this was not true for 2*L* > 2. This setting allowed us to assess the impact of increasing the number of fitness-related loci within the inversion beyond two while either keeping the total map distance of the inversion constant or increasing it (roughly linearly) with the number of fitness-related loci within the inversion.

We further assumed that mating was random within each local population, that is, the pool of *2N* gametes obtained after selection and recombination was randomly divided into *N* pairs, thus producing *N* offspring in each population (this corresponds to sampling without replacement). After reproduction, all adult individuals were replaced by their offspring, which were then treated as the next generation of adults.

As mentioned above, we studied two out of the five models analyzed by Feder and Nosil (2009), specifically their Model 2 that involves alleles that are locally favored in one habitat and neutral in the other, and Model 4 that involves universally beneficial alleles with negative fitness epistasis (*i.e.*, incompatibilities). In both models and in line with Feder and Nosil (2009), we considered two types of loci, denoted here by ***A***_*i*_ and ***B***_*i*_ for *i* = 1,2, …, *L* (the difference between the two types of loci is explained further below). Initially, all loci were bi-allelic: the two alleles at locus ***A***_*i*_ (***B***_*i*_) are denoted by *A*_*i*_ and *a*_*i*_ (that is, *B*_*i*_ and *b*_*i*_ for locus ***B***_*i*_). Furthermore, the alleles denoted by uppercase letters at each locus were initially fixed in Population 1, whereas the alternative alleles (denoted by lowercase letters) were initially fixed in Population 2.

Selection was implemented in the two models as follows. In Model 2, the alleles denoted by uppercase letters at loci ***A***_*i*_ (*i* = 1,2, …, *L*) were advantageous over the alternative alleles at these loci in the first population, whereas both allelic types at these loci were selectively neutral in the second population. The opposite was true for loci ***B***_*i*_ (*i* = 1,2, … , *L*): whereas at these loci both allelic types were selectively neutral in the first population, the alleles denoted by lowercase letters were advantageous over the alternative alleles in the second population. We considered two different models with respect to how loci ***A***_*i*_ and ***B***_*i*_ were ordered in the genomic region simulated. First, we considered the case where loci of the type ***A***_*i*_ were next to each other on the first half of the region (taking places 1,2, … , *L*), and loci of the type ***B***_*i*_ were on the second half of the region (thus, taking places *L* + 1, *L* + 2, … , 2*L*). We refer to this version of the model throughout as the *half-half array*. Second, we considered the case where loci of the two types were arranged consecutively (i.e. interdigitated), such that the order of loci was ***A***_1_, ***B***_1_, ***A***_2_, ***B***_2_, … , ***A***_*L*_, ***B***_*L*_. We refer to this version of the model as the *consecutive array*. These two versions of the model (relative to how the two types of loci are arranged in the inversion) present two extreme cases chosen here for convenience. Of course, real systems will present a continuum of possibilities between these two extremes.

We denoted selection coefficients per locus by *s*_2*L*_. For individuals that are homozygotes for the advantageous allele, heterozygotes, or homozygote for the disadvantageous allele (relative to the alternative allele), a fitness-related locus (see above) made a fitness contribution equal to 1 + *s*_2*L*_, 1 + *s*_2*L*_/2, and 1, respectively, with fitness being multiplicative across loci. Note that we conservatively scaled selection to avoid modeling the trivial case of extremely strong selection. To do so, we chose the selection coefficient per locus to depend on the number of loci to achieve the same maximal fitness when varying the number of loci within the inversion polymorphism. Denoting the maximal fitness by 1 + *s*, and recalling that only *L* loci were under selection in each population (whereas *L* loci were neutral), it follows that 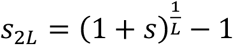, with *s*_2_ = *s*, as expected.

In Model 4 with universally beneficial alleles and negative epistasis, alleles *A*_*i*_ at loci ***A***_*i*_ were favored in both populations, whereas at loci ***B***_*i*_, the universally beneficial alleles were those denoted by lowercase letters (*i.e.*, ***b***_*i*_). As in the previous model, loci contributed to fitness in a multiplicative manner with the selection coefficient per locus (*s*_2*L*_) depending on the number of loci *2L* within the inversion polymorphism.

Genetic incompatibilities were modelled as occurring between universally favored alleles *A*_*i*_ and *b*_*i*_ at pairs of loci ***A***_*i*_ and ***B***_*i*_ (note the same index *i*). We accounted for incompatibilities slightly differently from Feder and Nosil (2009). Namely, for each individual we counted the number of incompatibilities at each pair of loci ***A***_*i*_ and ***B***_*i*_ as follows. (1) If an individual is a homozygote for allele *A*_*i*_ and a heterozygote at locus ***B***_*i*_, we accounted for two incompatibilities at this pair of loci; (2) for a homozygote for allele *A*_*i*_, and a homozygote for allele *b*_*i*_, we accounted for four incompatibilities for this pair of loci; (3) for an individual that is a heterozygote at both loci ***A***_*i*_ and ***B***_*i*_, we counted one incompatibility; and (4) for an individual that is a heterozygote at locus ***A***_*i*_ but a homozygote for allele *b*_*i*_ at locus ***B***_*i*_, we accounted for two incompatibilities at this pair of loci. All other allele combinations at these two pairs of loci gave zero incompatibilities. For each individual we thus computed the number of incompatibilities at each pair of loci ***A***_*i*_ and ***B***_*i*_ (*i* = 1,2, …, *L*), and summed them up to obtain the total number of incompatibilities (denoted by *n*_inc_) that the individual carried. To each individual that carried at least one incompatibility (i.e. *n*_inc_ ≥ 1), we assigned a fitness of 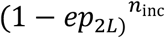, with *ep*_2*L*_ denoting the strength of the negative epistasis per incompatibility. Note that, because we assumed that incompatibilities may arise between pairs of loci, the maximum number of incompatibilities in the model with 2*L* loci is equal to 4*L*, and this is obtained when an individual is a homozygote for alleles *A*_*i*_ and for alleles *b*_*i*_ at all loci ***A***_*i*_ and ***B***_*i*_. To assure meaningful comparison between the models with different number of fitness-related loci within the inversion, we set *ep*_2*L*_ = 1 − (1 − *ep*)^1/*L*^, so that when 2*L* = 2, we have *ep*_2_ = *ep*. Note that our implementation of incompatibilities differs from that in Feder and Nosil (2009), where it was assumed that an individual carrying any combination of incompatible alleles has a fitness disadvantage equal to 1 − *ep* (independently of how many incompatibilities it has and of their linkage phase on chromosomes, as long as there is at least one). Similar to the model with locally favored and neutral alleles, here we also analyzed two different orderings of the loci ***A***_*i*_ and ***B***_*i*_, that is, we considered the *half-half array* ordering, and the *consecutive array* ordering (see above).

The parameter values we tested corresponded largely to those tested by Feder and Nosil (2009), but here we focused on a much larger rate of gene flux (by four orders of magnitude) than that used by them, and even larger in the *increasing-size* version of the models with more than two loci (see above). In addition, and unlike Feder and Nosil (2009), here we examined the effects of finite population size using populations of size *N* = 1,000 (but, for some parameter values, we also performed simulations with *N* = 5,000, *N* = 10,000, *N* = 20,000, and *N* = 100,000. Finally, we assessed the effect of increasing the number of loci within the inversion (2*L* = 2, 4, 6, 8, 10 or 20), while keeping the total selection strength constant (details explained above). For the full list of parameter values we used see Table I, as well as Tables S1–S2.

**Table I.** Parameters, their symbols, explanations, and values used in computer simulations.

| Symbol | Explanation | Values |
| --- | --- | --- |
| $N$ | Number of diploid individuals in each population | 1000, 5000, 10000, 20000, 100000 |
| $m$ | Per-individual per-generation migration probability (migration rate) | 0.01, 0.1 |
| $r$ | Per-gamete per-generation recombination probability between loci in the two locus models; with more loci ($2L$), the per-gamete per-generation recombination probability between a pair of consecutive loci is set to $r_{2L} = 1 - (1 - r)^{1/(2L-1)}$ (in the *conserved-size* version of the models), or to $r$ (in the *increasing-size* version of the models). | 0.1 |
| $r_{Inv}$ | Per-heterokaryotype per-generation recombination probability (gene flux) between the alternative chromosomal arrangements in the *conserved-size* version of the models; in the *increasing-size* version of the models, the total rate of gene flux is set to $r_{Inv} L$. | $0, 2 \cdot 10^{-4}$ |
| $s$ | Total selection strength | 0.1 |
| $2L$ | Number of loci within an inversion polymorphism | 2, 4, 6, 8 10, 20 |
| $ep$ | Strength of the negative epistasis | 0.1, 0.5, 0.95 |

In addition, for each model and each parameter set, we ran comparative simulations without any inversion polymorphism (*i.e.*, considering only collinear regions). In these simulations, we set the recombination rate between pairs of loci to be equal to the corresponding recombination rate in homokaryotypes in the model with inversion polymorphism. In comparison, Feder and Nosil (2009) assumed a recombination rate of 0.5 in their simulations of collinear regions.

In each simulation, we tracked the evolution of the allelic content of the inversion polymorphism for 100,000 generations after the populations come into secondary contact. The number of independent realizations we performed per parameter set was 200 (for *N* = 1,000, and *N* = 5,000), 40 (for *N* = 10,000, and *N* = 20,000), or 20 (for *N* = 100,000). During each simulation, we recorded (in steps of Δ*t* = 50 generations) the allele frequencies at each locus within the inversion polymorphism, the frequencies of alternative chromosomal arrangements in each population, as well as *F*_ST_ and *D*_*xy*_ at each locus. To facilitate comparisons between the simulation results corresponding to different parameter sets, we calculated the average time of sustained differentiation weighted by the differentiation (*T*_*w*_; hereafter referred to as the *average weighted time of differentiation*):

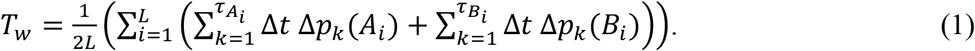

Here, the factor 1/(2*L*) serves to average over all fitness-related loci, and index *k* accounts for all sampling time points when the recorded allele-frequency difference Δ*p*_*k*_(*A*_*i*_) (or Δ*p*_*k*_(*B*_*i*_)), at a given locus ***A***_*i*_ (resp. ***B***_*i*_) was nonzero (here, Δ*p*_*k*_(*A*_*i*_) represents the frequency difference of allele *A*_*i*_ between Population 1 and Population 2; and similarly for Δ*p*_*k*_(*B*_*i*_)). The last such sampling point for locus ***A***_*i*_ (***B***_*i*_) is denoted 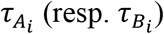. Note that in the absence of mutations, which we assumed, once the allele-frequency difference between the two populations becomes zero, it stays zero infinitely. However, we did not observe fixation in many cases we simulated because we ran simulations only up to a maximum of 100,000 generations after secondary contact. Therefore, the upper bound for *T*_*w*_ in our simulations was 100,000 generations, even if the actual average weighted time of differentiation would, in fact, be longer. Finally, Δ*t* in Eq. (1) stands for the time span between consecutive sampling points in our simulations (Δ*t* = 50).

We note that in the model without gene flux, due to our scaling of the strength of selection and of negative epistasis, the number of loci within the inversion does not play a role in the temporal dynamics of between-population differentiation. Consequently, all results for the cases without gene flux were obtained by averaging over the simulation results corresponding to different numbers of loci (except for *D*_*xy*_ and *F*_ST_, where we show the corresponding statistics for each locus obtained in a single randomly chosen stochastic realization of the model).

## Results

In Model 2 involving neutral and universally beneficial alleles, our simulations showed that when the fitness-related loci were captured by an inversion polymorphism (either with or without gene flux), the persistence time of differentiation weighted by the differentiation after secondary contact (*T*_w_; see Model and Methods) was longer than in the absence of inversion polymorphism: the difference between the two ranged from thousands to tens of thousands of generations (Figs. 1-2, and see Figs. S1-S10; but note that the actual persistence time of differentiation after secondary contact was larger than *T*_w_, due to the weighting we made when defining *T*_w_). This was true both when gene flux between the alternative arrangements was absent, as well when it was relatively high (in particular, four orders of magnitude higher than that assumed in Feder and Nosil, 2009). Notably, when gene flux was present, the weighted persistence time of differentiation increased rapidly with increasing number of fitness-related loci within the inversion. This was not the case in the model without inversions (although in this case a slight increase of *T*_w_ with increasing the number of loci occurred in the *consecutive-array*, *conserved-size* variation of the model; Fig. 1c-d). Our results, thus, showed that the advantage of inversions in maintaining population differentiation after secondary contact increased with increasing the number of fitness-related loci within the inversion, with *T*_w_ usually reaching a plateau after six, or more loci involved. This was true for all parameter values and variations of the model we considered, but the advantage was typically stronger for lower than for higher migration rates, and in the *conserved-size* than in the *increasing-size* variation of the model (compare Fig. 1 and Fig. 2). Our simulations showed that there were also some differences between the model variations with respect to the ordering (*half-half array* or *consecutive array*) of the two types of loci (***A***_*i*_ that carried locally beneficial alleles in Population 1, and ***B***_*i*_ that carried locally beneficial alleles in Population 2), but these differences were subtle.

**Figure 1.**
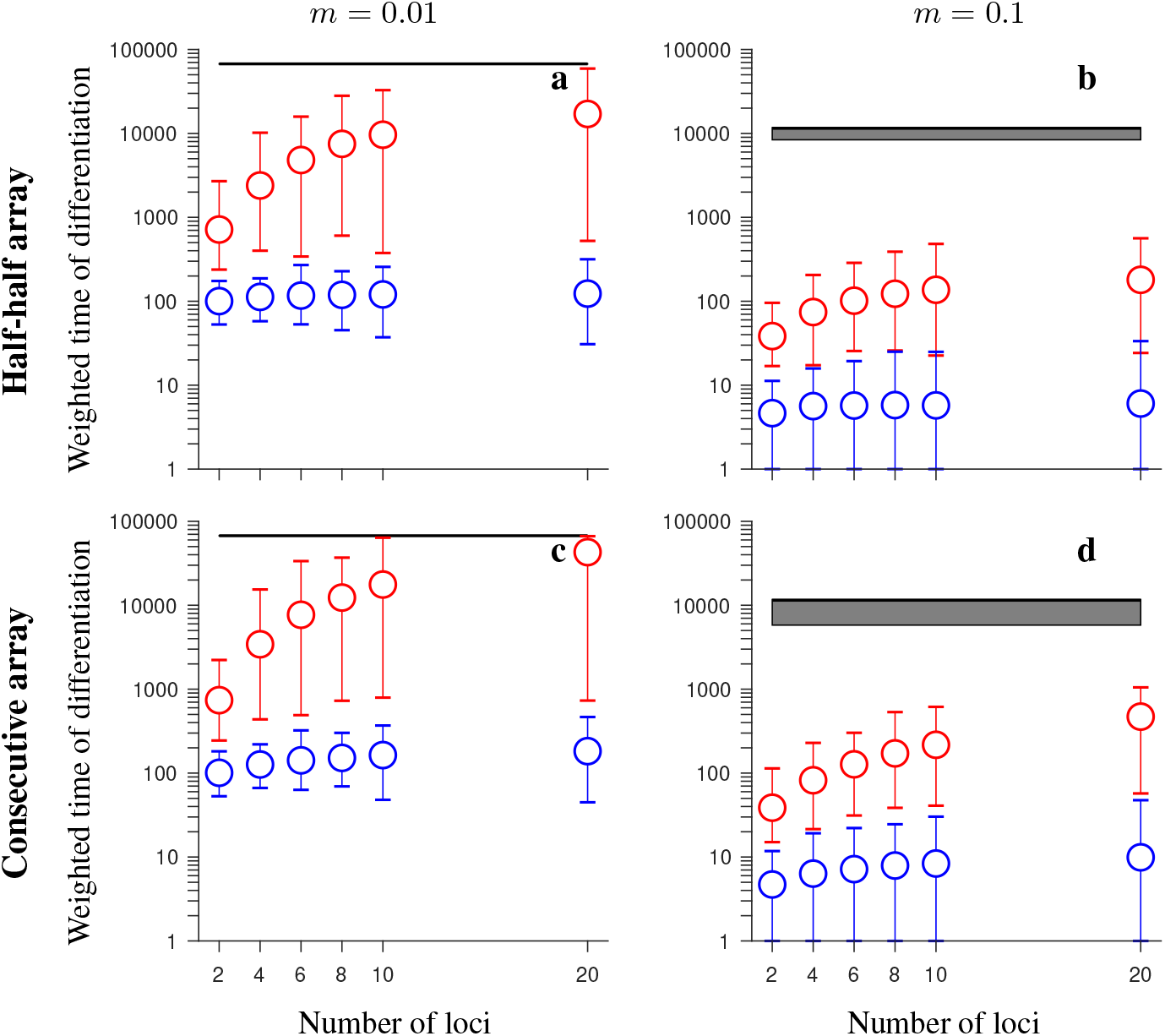
Simulation results for the *conserved-size* model involving locally favored and neutral alleles. The figure shows the weighted time of differentiation (*T*_*w*_) averaged over 200 independent realizations of the model, and over all loci within the region, as a function of the number of loci. Note the logarithmic scale on the vertical axis on this and all subsequent figures. Gene flux in heterokaryotypes is *r*_*Inv*_ = 2 · 10^−4^ (red) or *r*_*Inv*_ = 0 (black horizontal lines). Results for the model without inversions are shown in blue. The vertical lines around the symbols, and the grey regions (for *r*_Inv_ = 0) depict the range between the minimum and maximum values of *T*_*w*_ obtained in individual simulations. The panels differ by the migration rate (*m*) and by the ordering of the loci ***A***_*i*_ and ***B***_*i*_, as indicated in the figure. Remaining parameters: selection strength *s* = 0.1, number of individuals in each population *N* = 1,000, recombination rate in homokaryotypes *r* = 0.1. This figure shows that, despite gene flux, inversions maintain population differentiation after secondary contact for notably longer time periods than collinear regions, and this effect becomes stronger as the number of fitness-related loci increases.

**Figure 2.**
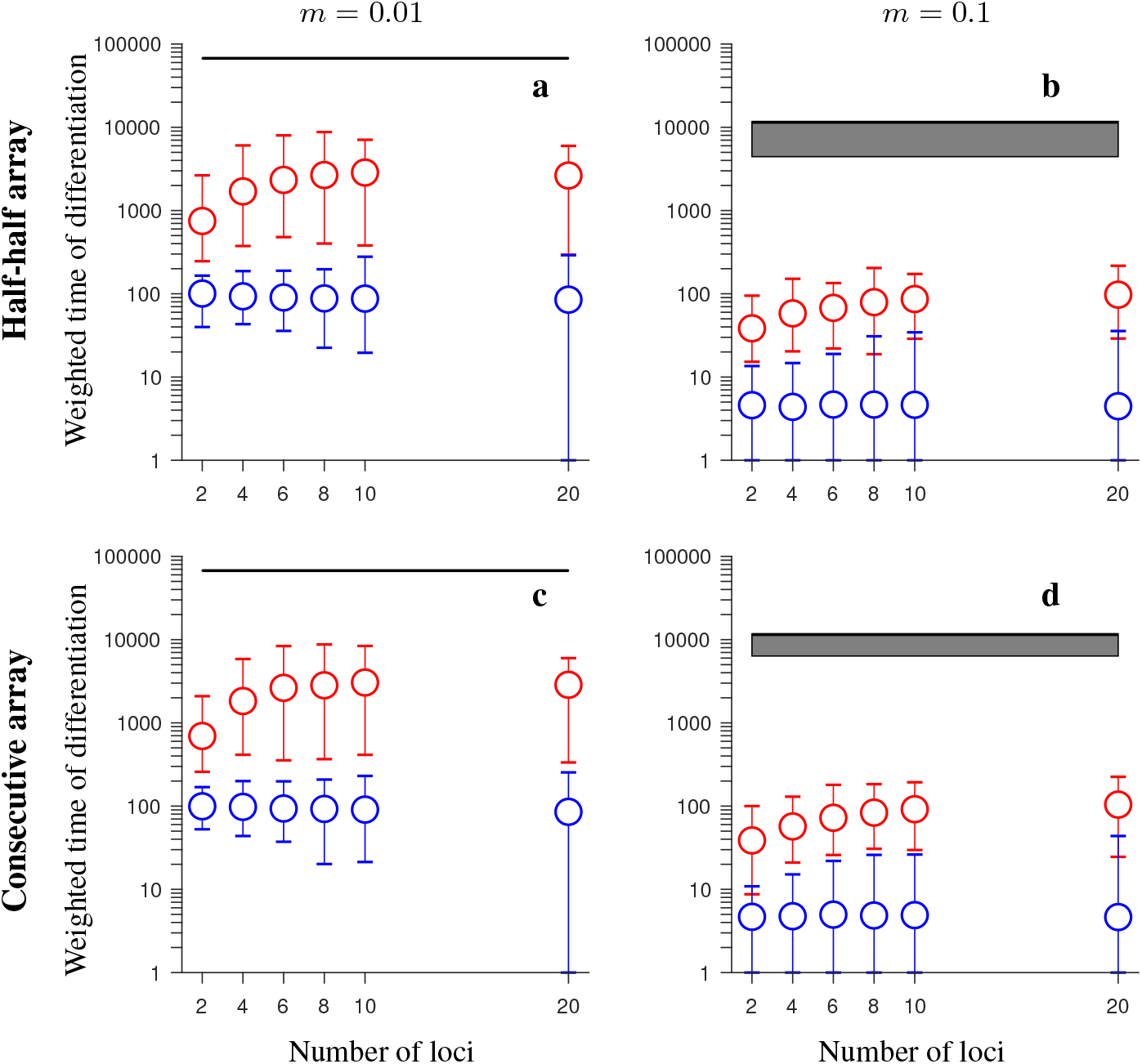
Same as in Fig. 1 but for the *increasing-size* version of the model with locally favored and neutral alleles (*i.e*., the total recombination distance within the region considered, as well as the total rate of gene flux, increase with increasing numbers of loci). This figure shows that inversions maintain population differentiation after secondary contact for notably longer time periods than collinear regions, and this effect becomes stronger as the number of fitness-related loci increases (although not as strongly as for the *conserved-size* model in Fig. 1), despite the fact that the rate of gene flux increases linearly with the number of loci.

Stronger and more persistent population differentiation after secondary contact in the presence of inversion was also observed when comparing *F*_ST_ (and *D*_*xy*_) patterns arising in individual realizations of the models with and without inversions (Figs. S2-S5, S7-S10). However, we found that *F*_ST_ (*D*_*xy*_) values were relatively low when the migration rate was high (i.e., of the order of the total selection strength) even in the presence of inversion polymorphism (panels d-f in Figs. S2-S5, S7-S10). Note that stochastic fluctuations between individual realizations were high (not shown).

In contrast to the model involving neutral and universally beneficial alleles, we found that differences in simulations involving genetic incompatibilities with and without inversions depended more strongly on the parameter values. When migration was weak in comparison to selection and negative epistasis was weak, inversions maintained population differentiation after secondary contact for longer than collinear regions, despite gene flux, and this difference was greater when the number of loci was larger (Figs. 3a, 4a, 5a, 6a). In the *conserved-size* version of the model, this effect was stronger in the *half-half array* than in the *consecutive-array* version of the model (compare Fig. 3a to Fig. 4a). In contrast, in the *increasing-size* version of the model, this effect was similar for the *half-half array* and the *consecutive array* versions of the model (compare Fig. 5a to Fig. 6a). Notably, the difference in persistence time of population differentiation in the presence versus in the absence of inversion polymorphism was higher in the *increasing-size* than in the *conserved-size* version of the model. This was because in the *increasing-size* model, the total recombination rate along the collinear region (model without any inversion polymorphism) increased with increasing numbers of loci within the region, and this significantly reduced the persistence time of population differentiation after secondary contact in the absence of inversions (down to 100 generations on average; Figs. 5a, 6a; blue circles). Note that in the *increasing-size* model, the total rate of gene flux also increased (linearly) with the number of loci. However, in this case, gene flux occurs only in heterokaryotypes and the frequency of heterokaryotypes after secondary contact was typically low for the parameters considered (Figs. S11a, S14a, S17a, S20a). Thus, for populations of modest to small sizes, this meant that the effective gene flux rate was still relatively rare despite being higher than in the *conserved-size* model.

**Figure 3.**
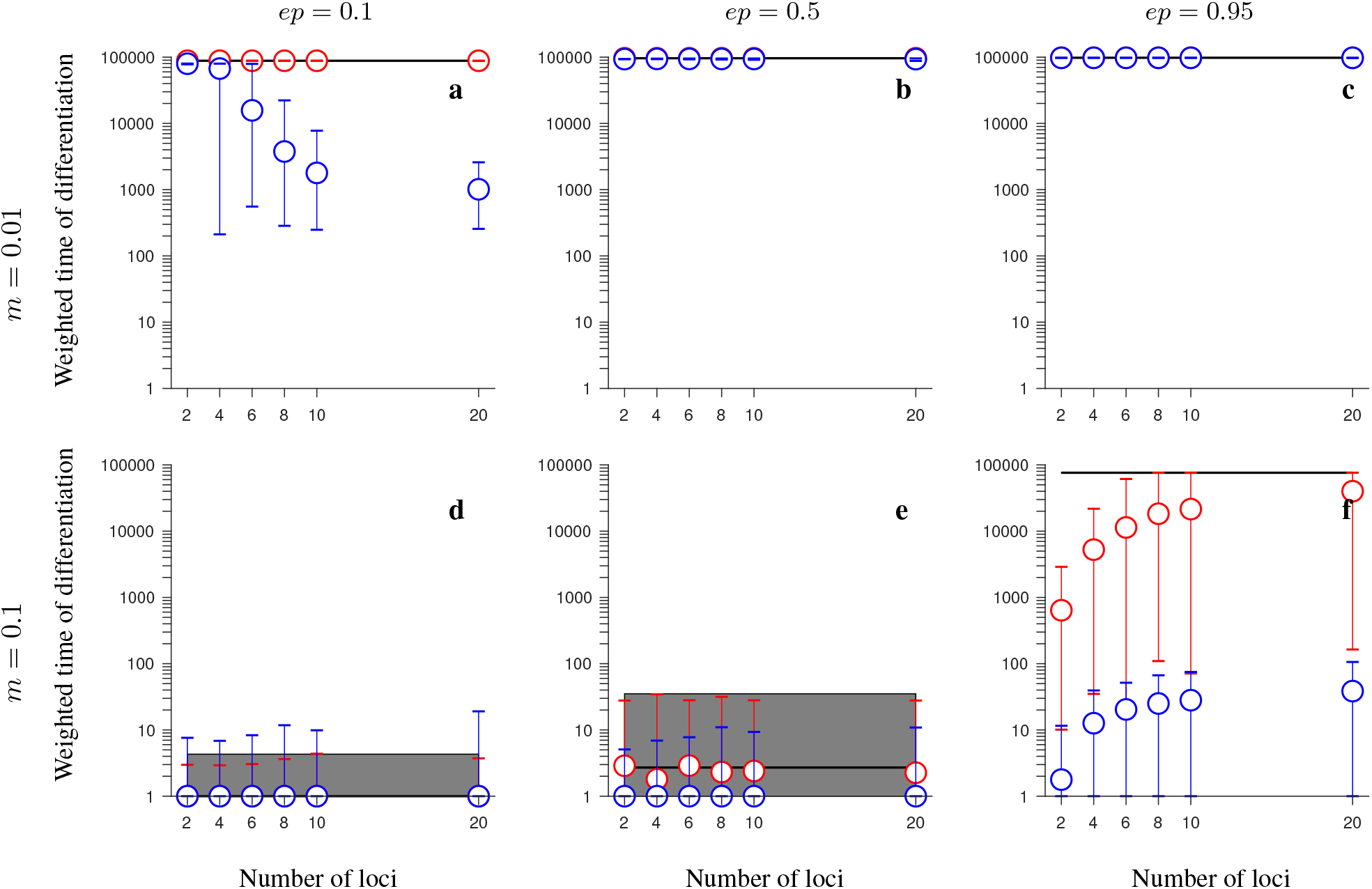
Simulation results for the *conserved-size*, *half-half array* model involving universally beneficial alleles with genetic incompatibilities. The figure shows the weighted time of differentiation (*T*_*w*_) averaged over 200 independent realizations and over all loci within the region, as a function of the number of loci. -Gene flux in heterokaryotypes is *r*_Inv_ = 2 · 10^−4^ (red) or *r*_Inv_ = 0 (black). Results for the model without inversions are shown in blue. Note that red and blue circles overlap in panels **b**-**d**. The vertical lines around the symbols, and the grey regions (for *r*_Inv_ = 0) depict the range between the minimum and maximum values of *T*_*w*_ obtained in individual simulations. The panels differ by the migration rate (*m*), and by the strength of negative epistasis (*ep*), as indicated in the figure. Remaining parameters: selection strength *s* = 0.1, number of individuals in each population *N* = 1,000, recombination rate in homokaryotypes *r* = 0.1. This figure shows that, when genetic incompatibilities and migration are weak, or when genetic incompatibilities and migration are strong, inversions (despite gene flux) maintain population differentiation after secondary contact for notably longer time periods than collinear regions, and this effect becomes stronger as the number of fitness-related loci increases.

**Figure 4.**
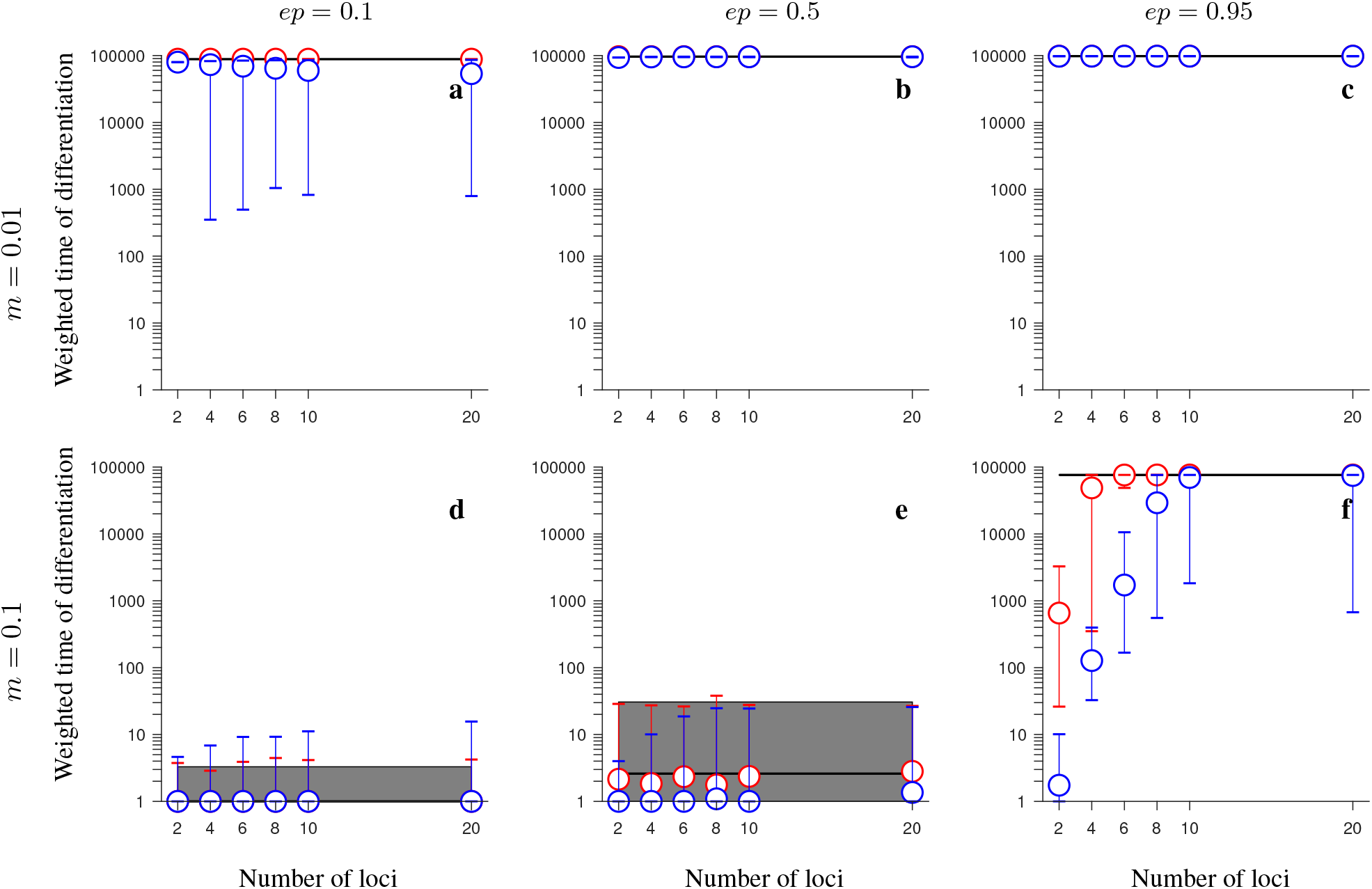
Same as in Fig. 3 but for the *consecutive-array* version of the *conserved-size* model involving universally beneficial alleles with genetic incompatibilities. Note that blue and red circles overlap in panels **b**-**d**. This figure shows that, despite gene flux, inversions may maintain population differentiation after secondary contact for notably longer time periods than collinear regions when genetic incompatibilities and migration are strong (panel f), but, unlike in the *half-half array* version of the model, this effect becomes weaker as the number of fitness-related loci increases, and it disappears when the number of loci is larger than 8.

**Figure 5.**
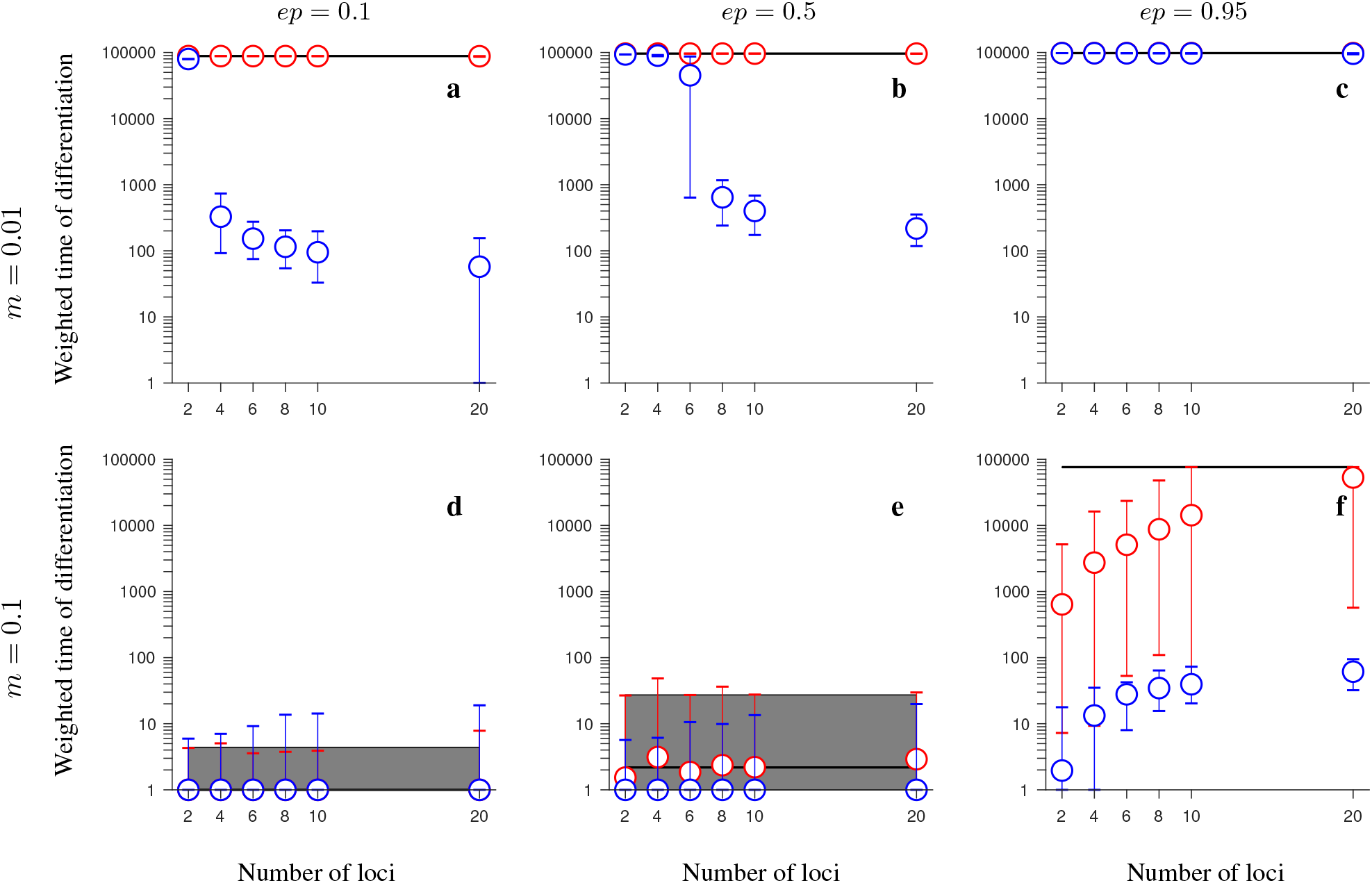
Same as in Fig. 3 but for the *increasing-size* version of the *half-half array* model involving universally beneficial alleles with genetic incompatibilities. Note that blue and red circles overlap in panels **c**, **d**. This figure shows that, when migration is weak and genetic incompatibilities are weak to modest, or when genetic incompatibilities and migration are strong, inversions maintain population differentiation after secondary contact for notably longer time periods than collinear regions, and this effect becomes stronger as the number of fitness-related loci increases, despite the fact that, in this version of the model, the rate of gene flux increases linearly with the number of loci.

**Figure 6.**
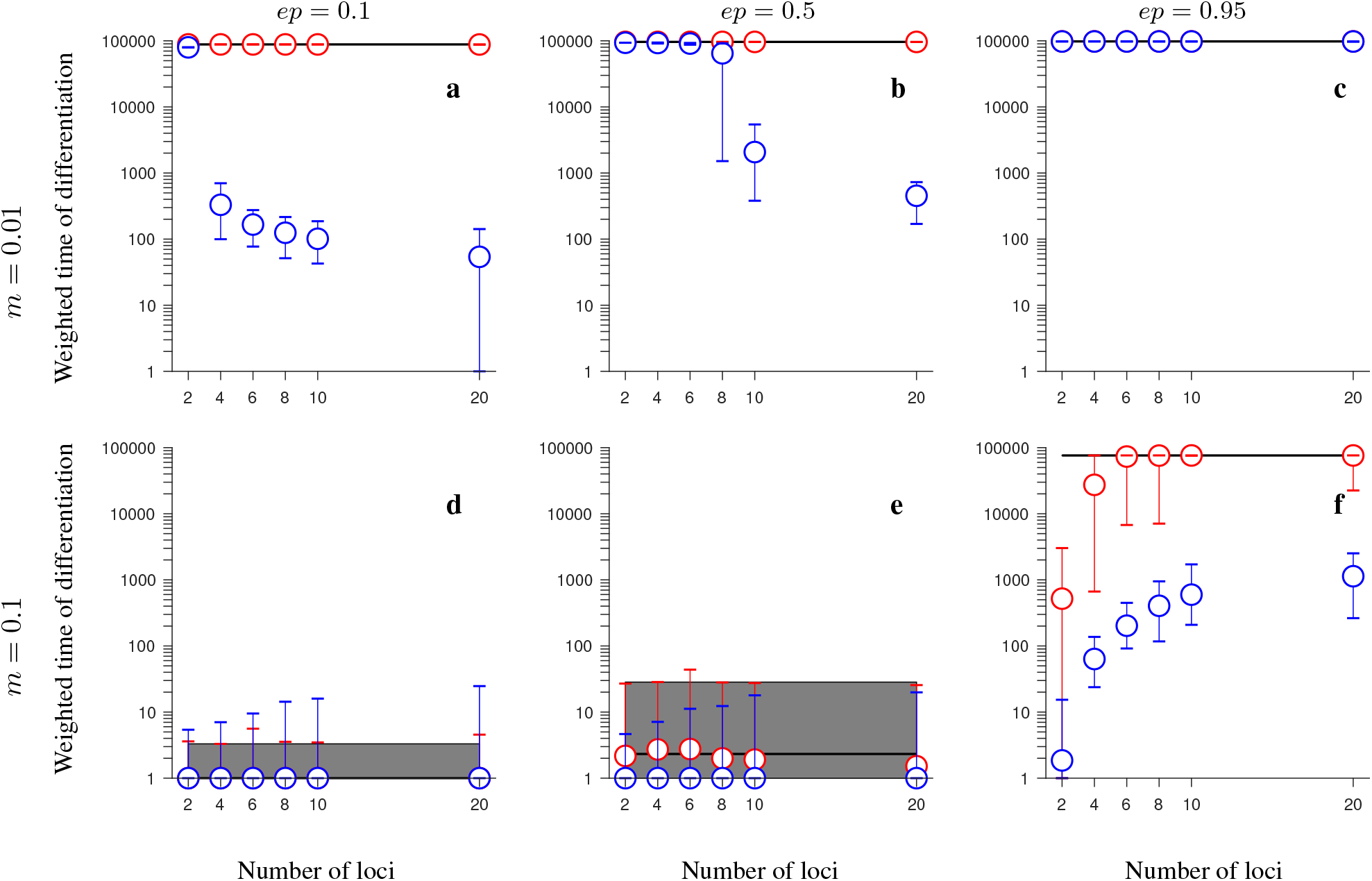
Same as in Fig. 3 but for the *increasing-size* version of the *consecutive-array* model involving universally beneficial alleles with genetic incompatibilities. Note that blue and red circles overlap in panels **c**, **d**. This figure shows that, when migration is weak and genetic incompatibilities are weak to modest, or when genetic incompatibilities and migration are strong, inversions maintain population differentiation after secondary contact for notably longer time periods than collinear regions, and this effect becomes stronger as the number of fitness-related loci increases, despite the fact that, in this version of the model, the rate of gene flux increases linearly with the number of loci. Note that this effect is slightly weaker than in the corresponding model with the *half-half array* ordering of loci.

For weak migration and intermediate negative epistasis, we found that strong population differentiation was maintained long after secondary contact (at least for 100,000 generations) independently of the number of loci in the model with inversion polymorphism and gene flux (red circles in Figs. 3b, 4b, 5b, 6d). The same was true without inversion polymorphism in the case of the *conserved-size* version of the model (blue circles in Figs. 3b, 4b). However, in the *increasing-size* version of the model without inversion polymorphism, population differentiation was lost more quickly when more fitness-related loci were involved (blue circles in Figs. 5b, 6b). As a result, population differentiation in this case was maintained for at least 500 times longer when loci were in the inversion than in the collinear region (compare red and blue circles in Figs. 5b, 6b; note that we say *at least*, because we ran simulations only up to 100,000 generations, and, for the parameters considered here, population differentiation was not lost during this time span in the model involving inversions with or without gene flux).

Conversely, for weak migration and very strong epistasis, population differentiation was maintained for at least 100,000 generations both in the model with and without inversions (Figs. 3c, 4c, 5c, 6c).

For strong migration and weak negative epistasis, population differentiation was lost quickly after secondary contact both with and without inversions (Figs. 3d, 4d, 5d, 6d), whereas for intermediate strength of negative epistasis, we found slightly longer persistence times of population differentiation in the case with inversions (with or without gene flux) than in the case without inversions, but this effect was mild (Figs. 3e, 4e, 5e, 6e).

Finally, when both migration and negative epistasis were strong, the persistence time of differentiation after secondary contact was, in most cases, by several orders of magnitude longer in the case with inversions (despite gene flux) than without inversions (Figs. 3f, 4f, 5f, 6f). This effect was stronger when the number of fitness-related loci was larger, except in the *consecutive-array* version of the model: in this case, in the *conserved-size* model without inversions, the persistence time of differentiation increased with increasing the number of loci (blue circles in Fig. 4f), reaching the maximum persistence time (within the time span we simulated) for ten loci. In the *consecutive-array*, *increasing-size* version of the model without inversions, the persistence time of differentiation also increased with increasing numbers of loci, but this increase was slower than in the *conserved-size* version of the model. For twenty loci, the persistence time of differentiation was shorter by two orders of magnitude in the case without inversions than in the case with inversions and gene flux (Fig. 6f). Note that the observed increase of persistence time of differentiation with increasing numbers of loci in the model without inversions in the case of high migration was opposite to what happened in the case of low migration (compare, for example, blue circles in Fig. 5a to blue circles in Fig. 5f). This was because in the case of high migration, the timescale of recombination between consecutive loci was either less than the migration timescale (in the *conserved-size* model), or the two timescales were similar (in the *increasing-size* model); and with increasing numbers of loci, the number of recombination events needed to purge incompatibilities increased. As a consequence, the persistence time of differentiation after secondary contact in the model without inversions increased with increasing numbers of loci in the high-migration case, whereas the opposite was true in the weak-migration case of the *increasing-size* model (where the timescale of recombination between consecutive loci was less than the migration timescale).

The patterns described for the model involving universally beneficial alleles with genetic incompatibilities were also reflected in *F*_ST_ (and *D*_*xy*_) patterns from individual realizations of the model (Figs. S12-S13, S15-S16, S18-S19, S21-S22). As in the model with locally beneficial and neutral alleles, stochastic fluctuations between individual realizations were high (not shown).

The findings outlined above were retained when considering populations of larger size (*N* = 5,000; Figs. S23-S24), but the advantage of an inversion polymorphism in maintaining population differentiation after secondary contact was slightly less than when *N* = 1,000 (compare Fig. 1a-b to Fig. S23a-b, and Fig. 3 to Fig. S24).

To further assess how population size impacts on the advantage of inversions over collinear regions in maintaining population differentiation after secondary contact, we performed simulations with population sizes of up to 100,000 individuals for a subset of parameter values, focusing on cases with 20 loci (Figs. 7-10). We found that the advantage of inversions over collinear regions was either fully retained when we increased the population size (Fig. 8a-d, 9a, 10a, 10c-d) or was less but still noticeable. However, in the *consecutive-array*, *conserved-size* version of the model involving universally beneficial alleles with genetic incompatibilities, population differentiation was strong and maintained throughout the entire simulated time period after secondary contact (100,000 generations) either with or without inversions (Fig. 9c-d).

**Figure 7.**
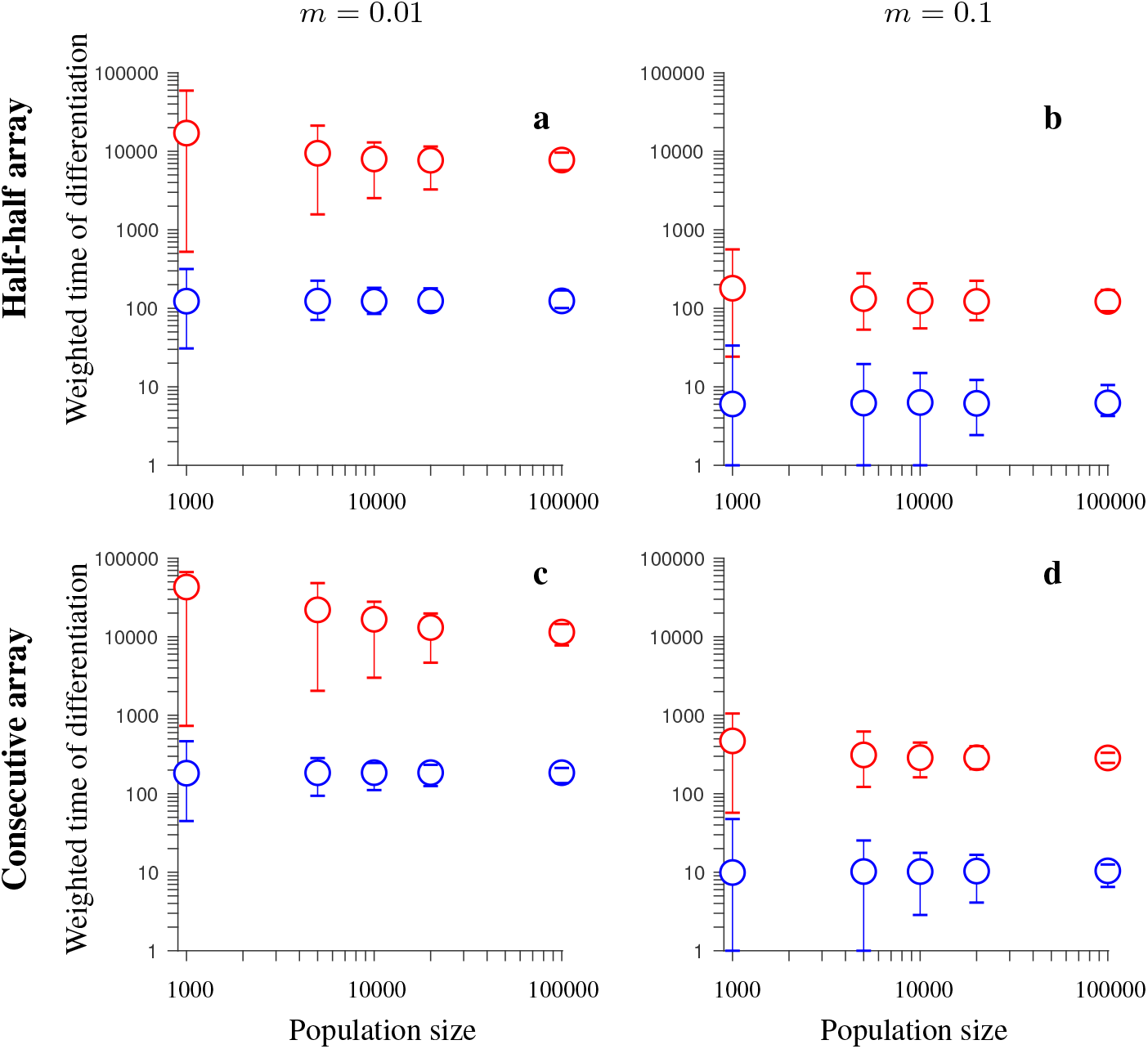
Simulation results for the *conserved-size* model involving locally favored and neutral alleles arranged at 20 loci. The figure shows the weighted time of differentiation (*T*_*w*_) averaged over independent realizations of the model, and over all loci within the region, as a function of the population size. Note the logarithmic scale on the vertical and horizontal axis. Gene flux in heterokaryotypes is *r*_*Inv*_ = 2 · 10^−4^ (red). Results for the model without inversions are shown in blue. The vertical lines around the symbols depict the range between the minimum and maximum values of *T*_*w*_ obtained in individual simulations. The panels differ by the migration rate (*m*) and by the ordering of the loci ***A***_*i*_ and ***B***_*i*_, as indicated in the figure. Remaining parameters: selection strength *s* = 0.1, recombination rate in homokaryotypes *r* = 0.1, number of independent realizations of the model is 200 (for *N* = 1,000, and *N* = 5,000), 40 (for *N* = 10,000 and *N* = 20,000), or 20 (for *N* = 100,000). This figure shows that the advantage of inversions (with gene flux) over collinear regions in maintaining population differentiation after secondary contact decreases with increasing the population size *N*. However, even for population sizes of *N* = 100,000, inversions (despite gene flux) maintain population differentiation after secondary contact for notably longer time periods than collinear regions.

**Figure 8.**
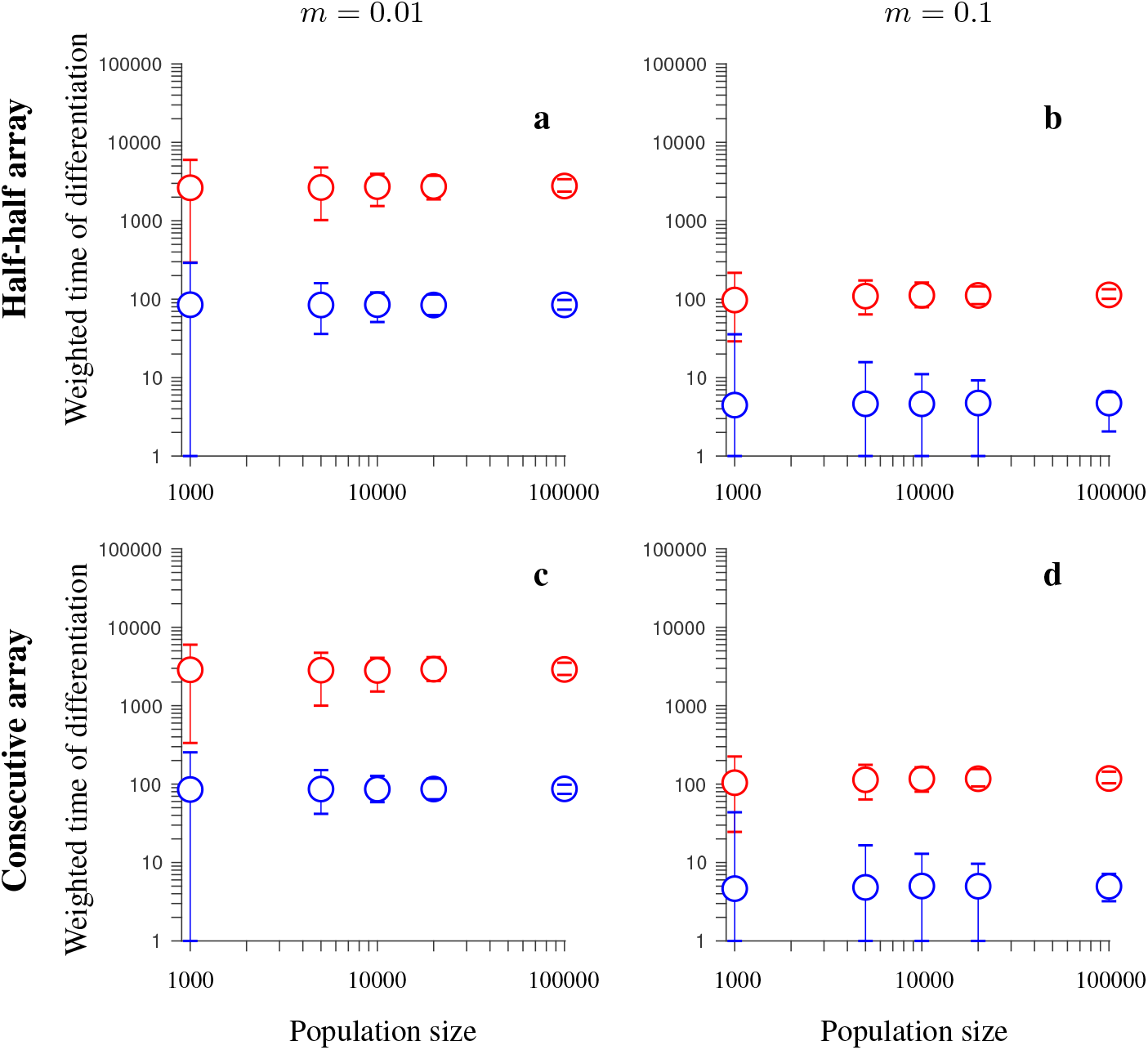
Same as in Fig. 7 but for the *increasing-size* version of the model with locally favored and neutral alleles (i.e., the total recombination distance within the region considered, as well as the total rate of gene flux increase with increasing the number of loci). This figure shows that the advantage of inversions (with gene flux) over collinear regions in maintaining population differentiation after secondary contact is roughly independent of the population size *N*, and it is notable even for population sizes of *N* = 100,000.

**Figure 9.**
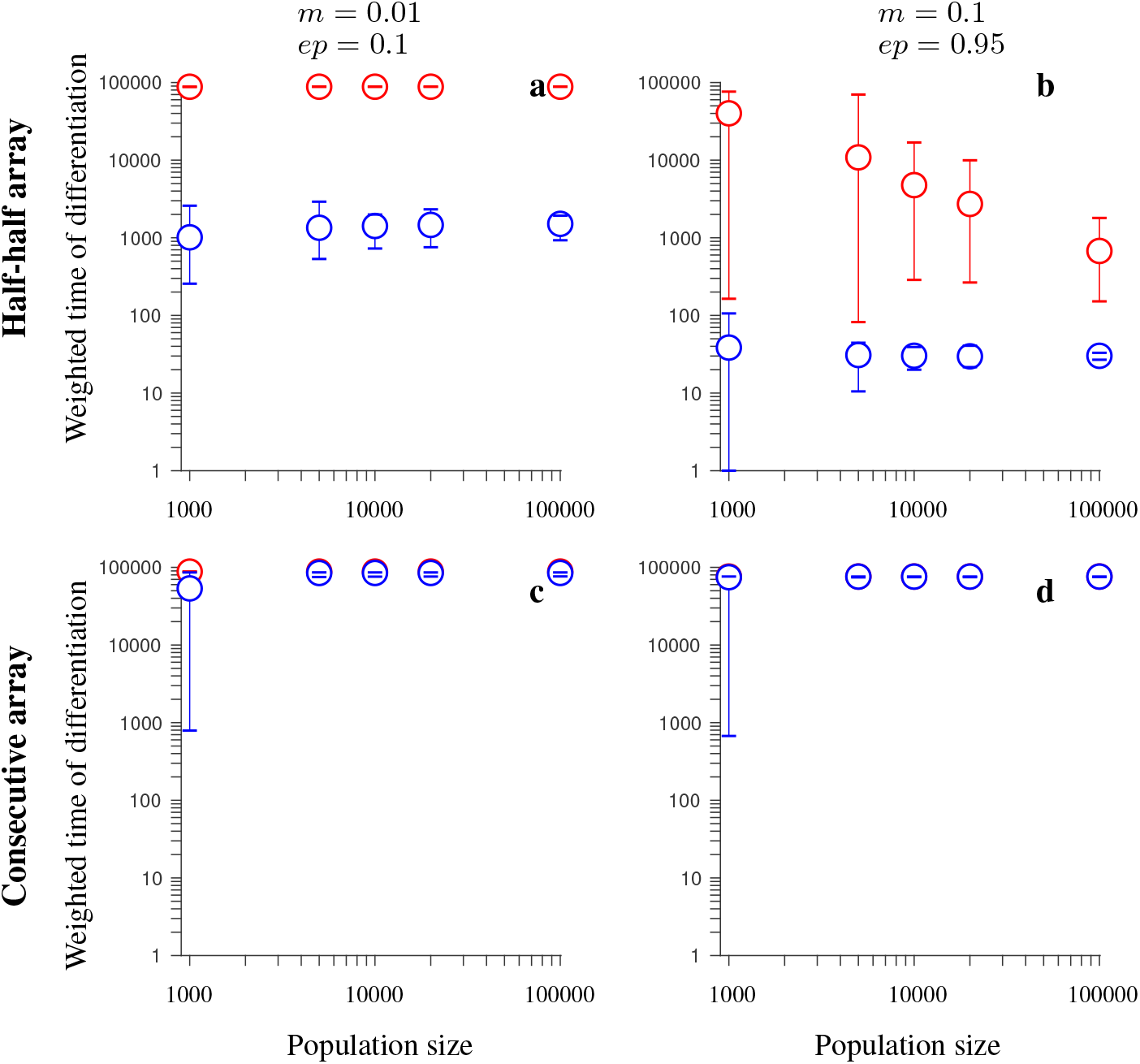
Same as in Fig. 7 but for the *conserved-size* version of the model involving universally beneficial alleles with genetic incompatibilities. Note that blue and red circles overlap in panels **c, d**. This figure shows that the advantage of inversions (with gene flux) over collinear regions in maintaining population differentiation after secondary contact is either roughly independent of the population size, *N* (panel a), or it decreases with increasing *N*, but it is notable even for population sizes of *N* = 100,000 (panel b), or that differentiation is maintained throughout the whole time span simulated both in the model with and without inversions (panels c, d).

**Figure 10.**
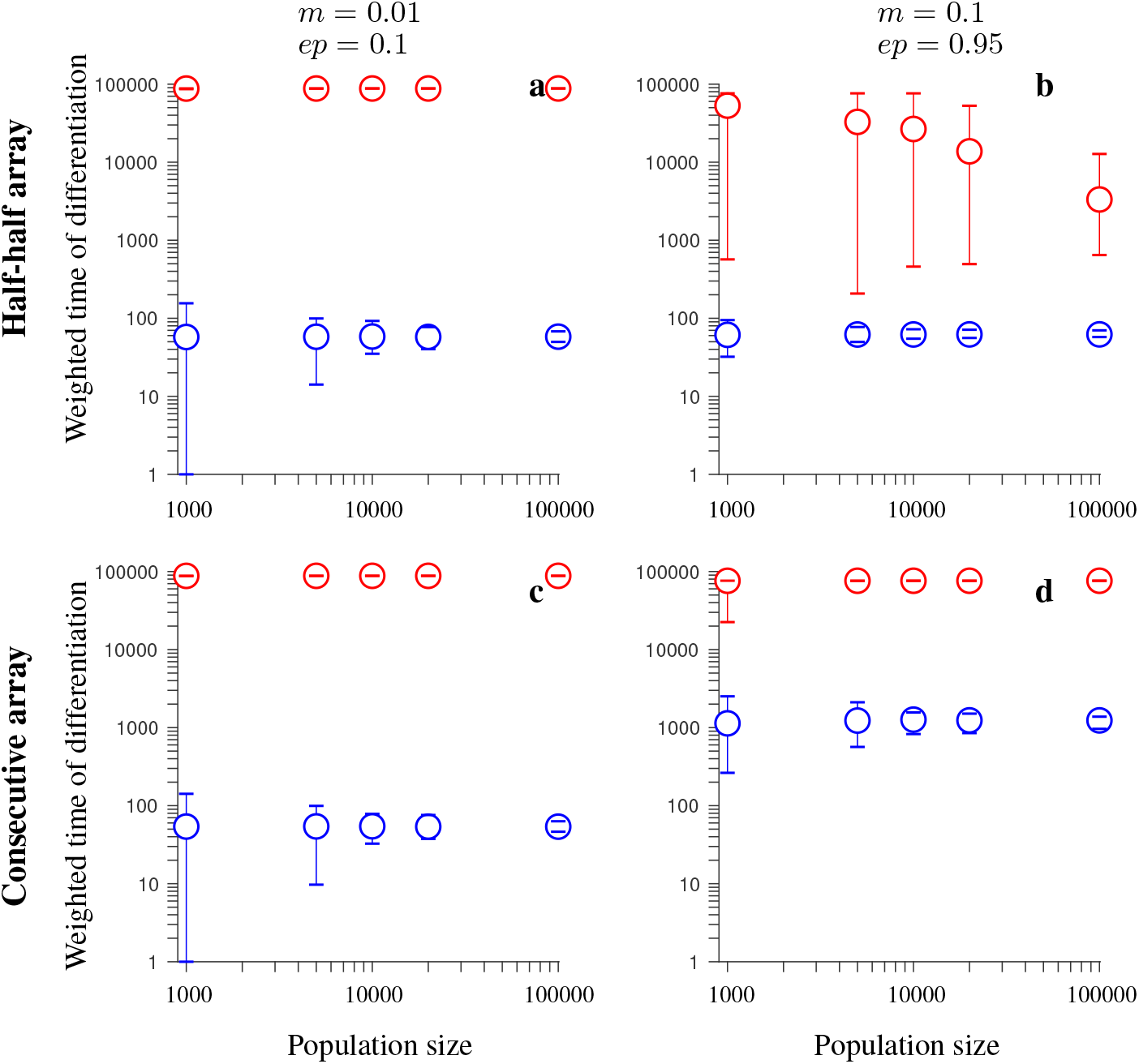
Same as in Fig. 9 but for the *increasing-size* version of the model involving universally beneficial alleles with genetic incompatibilities. This figure shows that the advantage of inversions (with gene flux) over collinear regions in maintaining population differentiation after secondary contact is either roughly independent of the population size, *N* (panels a, c, d), or it decreases with increasing *N*, but it is notable even for population sizes of *N* = 100,000 (panel b).

## Discussion

Earlier studies suggested that chromosomal inversions are semi-permeable barriers to gene flow, especially due to genetic exchange between the alternative arrangements by gene flux (Guerrero *et al.* 2012; Korunes and Noor 2016; Korunes and Noor 2019). Theoretical work by Feder and Nosil (2009) suggested that in the absence of strong local adaptation, population differentiation may tend to be eroded by gene flux and that this can occur quickly after secondary contact even with low gene flux, unless strong selection, and weak migration are at work. These results were difficult to reconcile with the repeated empirical observation of stronger population differentiation within regions polymorphic for inversions across different organisms (e.g. Kulathinal *et al.* 2009; Jones *et al.* 2012). However, Feder and Nosil (2009) based their results on models involving infinitely large populations and focused mainly on situations in which just a few loci reside within the inversion.

Here, we extended the analysis of Feder and Nosil (2009) for two models: one involving locally favored and neutral alleles (Model 2 in that study, also known as the conditional neutrality model), and the other involving universally beneficial alleles with negative epistasis (Model 4 in that study). We found that, in situations with finite population sizes and/or multiple loci, inversion polymorphism can, relative to collinear regions, delay the erosion of genetic differentiation between populations for tens of thousands of generations after secondary contact. Our findings differ from Feder and Nosil (2009) for three reasons. First, finite population sizes introduce a waiting time for gene-flux events to occur. This waiting time depends on the population size and the migration-selection quasi-equilibrium that is reached in each case, but it is in general longer in smaller than in larger populations. Second, even when gene flux does occur, any genotype it gives rise to must escape loss by drift. Thus, given that the probability of stochastic loss increases with decreasing effective population size, it follows that favored genotypes are more likely to be lost by genetic drift in smaller than in larger populations, thereby potentially delaying the elimination of population divergence. However, we note that this effect may be counteracted to some degree by a shorter fixation time of a genotype in smaller than in larger populations. Third, distributing selection across many loci in inversions, as opposed to being concentrated on a couple of genes, increases genetic interference, requiring a larger number of *favorable* gene flux events towards forming favorable genotypes, thereby decreasing the effectiveness of selection (on a haplotype level). This results in populations residing in a quasi-steady migration-selection equilibrium for a long period of time after secondary contact even for gene flux values as high as 2 · 10^−4^ per gamete per generation.

The rate of gene flux tested here is close to the upper bounds of the corresponding rates inferred from empirical data (between 10^−4^ and 10^−6^ converted sites/genome/generation across taxa (Korunes & Noor 2017). Recent data from the *D. pseudoobscura*-*D. persimilis* system imply that double crossover rates, although lower than in homokaryotypes or collinear regions, are non-negligible in heterokaryotypes (10^−4^) (Stevison *et al.* 2011). Gene conversion rates between 10^−5^ and 2.5 · 10^−5^ involving tracts of DNA sequences between 200 and 400bp long have also been observed for these species (Korunes and Noor 2019). While double crossovers occur mainly in the central parts of large inversions due to interference (Navarro *et al.* 1997a; Fuller *et al.* 2019), gene conversion rates seem to be similar across the entire inverted regions (without interference), including near the breakpoints (Korunes and Noor 2016; Crown *et al.* 2018).

Regardless of the mechanism of gene flux, our results show that inversions can delay population fusion after secondary contact for a longer period of time when they contain a larger number of loci under selection or involved in generating hybrid incompatibilities. However, the positional effects of the different mechanisms of gene flux, considering also interference between crossover events, need to be modelled in more detail. The previous deterministic analysis by Feder and Nosil (2009) suggested that spreading selection across greater number of loci in inversions could delay homogenization due to the longer time it takes for genes with lower per-allele selection strength to spread between populations. Our results advance understanding of the process by showing that finite population size and interference among linked loci can amplify the effects of multiple versus a few loci in curtailing the homogenization of rearranged regions of the genome following secondary contact.

We caution, however, that the difference between many versus a few loci we report here is valid when: (1) gene flux scaled by the effective population size is large enough (but not too large) to create a sufficient number of chromosomes possessing all favorable alleles to ensure homogenization; (2) per-allele selection strength scaled by population size is low enough so that rearrangements containing a small number of alleles under selection (following gene flux) will predominantly be lost; (3) effective population sizes are small to intermediate, with strong dependence on the details of the model and on parameter values; in some cases, the advantage of an inversion harboring many loci over collinear regions is retained even for populations with 100,000 individuals; and (4) favorable alleles in one population are neutral in the other population, or universally favorable alleles across populations cause incompatibilities in hybrids. We note that when fitness tradeoffs exist such that alleles beneficial in one habitat are detrimental in the other, inversions might still retard the genetic homogenization of finite populations but the effects of concentrating selection on a few versus many loci may differ. However, we leave the exploration of this model for future work. The extent to which the above conditions hold in nature will therefore have a significant bearing on the degree to which multiple loci prolong inversion differentiation following secondary contact. Finally, if rather than keeping the total strength of local adaptation or genetic incompatibility/negative epistasis constant and spreading it evenly over loci when additional genes are added, selection instead becomes stronger, the parameter space within which rearrangements will have a relevant role in population differentiation will vary drastically and remains to be explored in more detail.

One limitation of our study shared with earlier work is that we considered a two-population system because such a simple scenario is convenient for modelling. However, if testable quantitative predictions applicable to empirical systems are to be obtained, it will be necessary to use spatially-explicit models involving many local populations (Flaxman et al. 2012) with a spatial and/or temporal selection gradient (e.g. mimicking hybrid zones). Such models would provide a significant step towards understanding observations in many hybrid zones where genetic regions harboring inversion polymorphisms have been detected (*e.g.*, Rieseberg *et al.* 1999; Ayala *et al.* 2012; Westram *et al.* 2018; Faria *et al.* 2019a; Faria *et al.* 2019b). In addition, it would be appealing to model inversions that harbor alleles under divergent selection together with alleles that are universally beneficial (or deleterious), and/or alleles involved in incompatibilities, comparing cases with diffuse versus concentrated genetic architectures.

## Conclusions

Our results support the hypothesis that inversions may be a strong mechanism contributing to maintenance of population differentiation, and potentially speciation, when divergence has a polygenic basis (Feder *et al.* 2005; Kirkpatrick and Barton 2006). Crucially, by maintaining population differences after secondary contact, inversions can act as a partial barrier to gene flow between populations, thus allowing for a longer period of time compared to collinear gene regions during which populations may accumulate additional differences including incompatibilities. This may facilitate the speciation process to progress towards completion through other mechanisms, such as reinforcement (Noor *et al.* 2001; Butlin and Smadja 2018). Note that we do not claim that inversion polymorphisms are impermeable barriers to gene flow. Rather, unless new inversions arise or divergence by new mutations occurs, populations will eventually homogenize in the face of gene flow and in the absence of divergent selection. However, for many parameter sets we tested, homogenization can be delayed for tens of thousands of generations after secondary contact, and this delay can allow for additional barriers to accumulate.

Besides suggesting that chromosomal rearrangements have a role in maintaining population differences after secondary contact, previous theoretical work showed that they can also facilitate adaptive population differentiation in cases of primary contact, resulting in the emergence of clustered genomic architectures (Yeaman 2013). Moreover, reduced recombination rates within heterokaryotypes containing multiple weakly diverged loci will be an effective mechanism protecting such loci from stochastic loss (Feder et al. 2014; Rafajlović *et al.* 2016), thereby facilitating divergence with gene flow. Overall, these results, together with those presented here, suggest an important role of inversions in population divergence and speciation, even in the face of gene flux.

## Author contributions

AN and RF conceived the study. AN, RF and MR planned the simulations. JR coded and implemented initial simulations. MR coded and implemented the final simulations. MR analyzed the results. RF, MR and AN wrote the first draft. All authors contributed to the final version of the manuscript.

## Acknowledgments

The authors are indebted to Mohamed A.F. Noor and Roger K. Butlin for valuable comments on the manuscript. This study was supported by the European Regional Development Fund (FCOMP-01-0124-FEDER-014272), FCT – Foundation for Science and Technology (PTDC/BIA-EVF/113805/2009), Ministerio de Ciencia e Innovación, Spain (BFU2015-68649-P and PGC2018-101927-B-I00, MINECO/FEDER, UE), by the Spanish National Institute of Bioinformatics (PT17/0009/0020), by “Unidad de Excelencia María de Maeztu”, funded by the MINECO (ref: MDM-2014-0370). RF was financed by FCT (SFRH/BPD/89313/2012) and by the European Union’s Horizon 2020 research and innovation programme, under the Marie Sklodowska-Curie grant agreement number 706376. MR was funded by the Hasselblad Foundation (Grant for Female Scientists), European Research Council and the Swedish Research Councils VR and Formas (Linnaeus grant to the Centre for Marine Evolutionary Biology), and by additional grant from Formas (to MR). The simulations were performed on resources at Chalmers Centre for Computational Science and Engineering (C3SE) provided by the Swedish National Infrastructure for Computing (SNIC). JLF was funded by support from the National Science Foundation and United States Department of Agriculture NIFA program.

## Data Accessibility Statement

There is no empirical data to be archived in this manuscript.

## Supplementary Tables and Figures

**Table S1.**
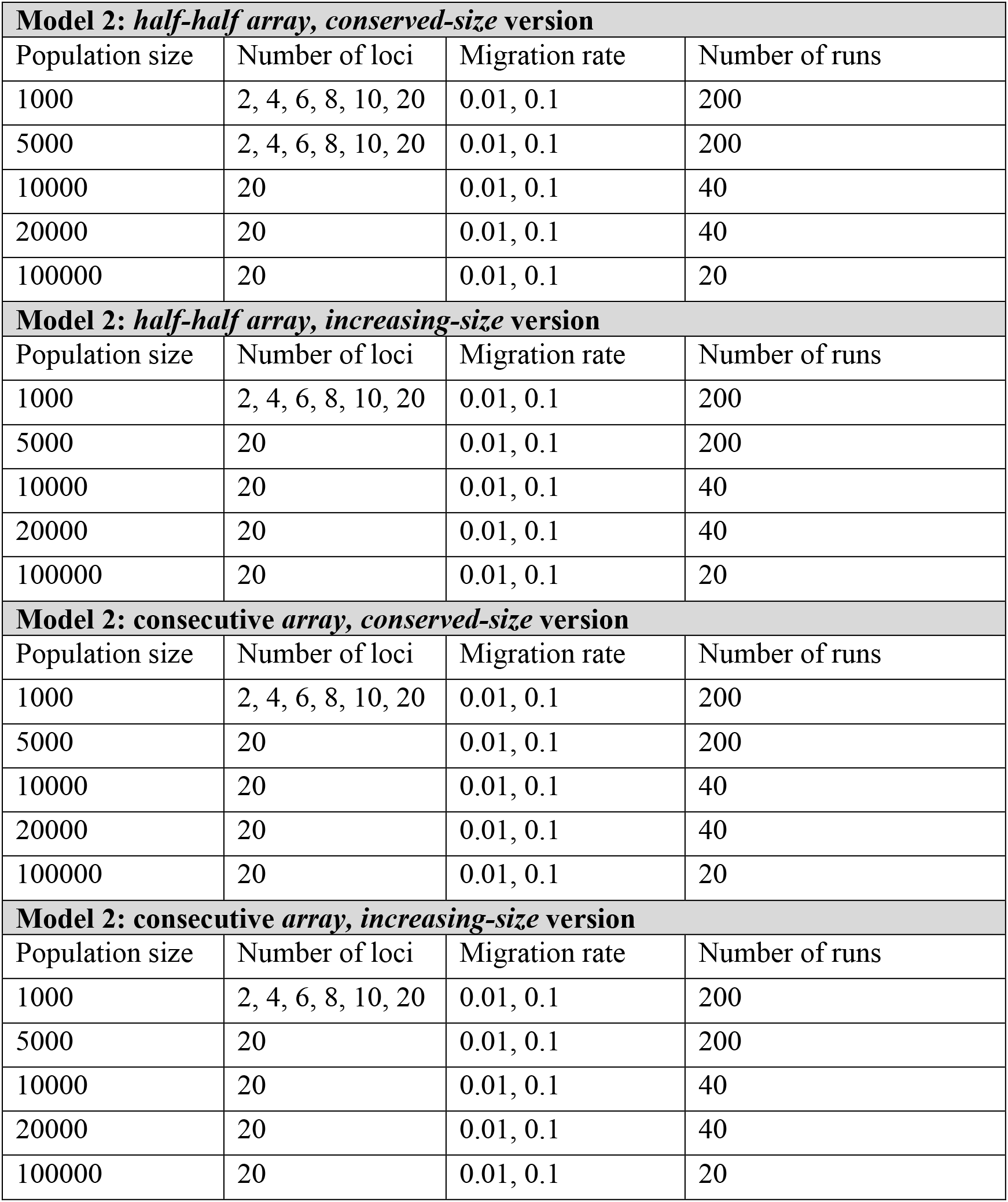
List of simulated versions of Model 2, and the corresponding combinations of parameter values used in the simulations. In all cases, selection strength was *s* = 0.1, and recombination rate was *r* = 0.1. In simulations with inversions, gene flux was either *r*_Inv_ = 0, or *r*_Inv_ = 2 · 10^−4^. For the explanation of the model, and parameters, refer to the main text and Table 1.

**Table S2.** List of simulated versions of Model 4, and the corresponding combinations of parameter values used in the simulations. In all cases, selection strength was *s* = 0.1, and recombination rate was *r* = 0.1. In simulations with inversions, gene flux was either *r*_Inv_ = 0, or *r*_Inv_ = 2 · 10^−4^. Note that, in some cases, only some parameter combinations for the migration rate and strength of epistasis were used. In these cases, the corresponding parameter combinations are denoted in the table by * or by **. For the explanation of the model, and parameters, refer to the main text and Table 1.

| **Model 4: half-half array, conserved-size version** |  |  |  |  |
| --- | --- | --- | --- | --- |
| Population size | Number of loci | Migration rate | Strength of epistasis | Number of runs |
| 1000 | 2, 4, 6, 8, 10, 20 | 0.01, 0.1 | 0.1, 0.5, 0.95 | 200 |
| 5000 | 2, 4, 6, 8, 10, 20 | 0.01, 0.1 | 0.1, 0.5, 0.95 | 200 |
| 10000 | 20 | 0.01*, 0.1** | 0.1*, 0.95** | 40 |
| 20000 | 20 | 0.01*, 0.1** | 0.1*, 0.95** | 40 |
| 100000 | 20 | 0.01*, 0.1** | 0.1*, 0.95** | 20 |
| **Model 4: half-half array, increasing-size version** |  |  |  |  |
| Population size | Number of loci | Migration rate | Strength of epistasis | Number of runs |
| 1000 | 2, 4, 6, 8, 10, 20 | 0.01, 0.1 | 0.1, 0.5, 0.95 | 200 |
| 5000 | 20 | 0.01*, 0.1** | 0.1*, 0.95** | 200 |
| 10000 | 20 | 0.01*, 0.1** | 0.1*, 0.95** | 40 |
| 20000 | 20 | 0.01*, 0.1** | 0.1*, 0.95** | 40 |
| 100000 | 20 | 0.01*, 0.1** | 0.1*, 0.95** | 20 |
| **Model 4: consecutive array, conserved-size version** |  |  |  |  |
| Population size | Number of loci | Migration rate | Strength of epistasis | Number of runs |
| 1000 | 2, 4, 6, 8, 10, 20 | 0.01, 0.1 | 0.1, 0.5, 0.95 | 200 |
| 5000 | 20 | 0.01*, 0.1** | 0.1*, 0.95** | 200 |
| 10000 | 20 | 0.01*, 0.1** | 0.1*, 0.95** | 40 |
| 20000 | 20 | 0.01*, 0.1** | 0.1*, 0.95** | 40 |
| 100000 | 20 | 0.01*, 0.1** | 0.1*, 0.95** | 20 |
| **Model 4: consecutive array, increasing-size version** |  |  |  |  |
| Population size | Number of loci | Migration rate | Strength of epistasis | Number of runs |
| 1000 | 2, 4, 6, 8, 10, 20 | 0.01, 0.1 | 0.1, 0.5, 0.95 | 200 |
| 5000 | 20 | 0.01*, 0.1** | 0.1*, 0.95** | 200 |
| 10000 | 20 | 0.01*, 0.1** | 0.1*, 0.95** | 40 |
| 20000 | 20 | 0.01*, 0.1** | 0.1*, 0.95** | 40 |
| 100000 | 20 | 0.01*, 0.1** | 0.1*, 0.95** | 20 |

**Figure S1.**
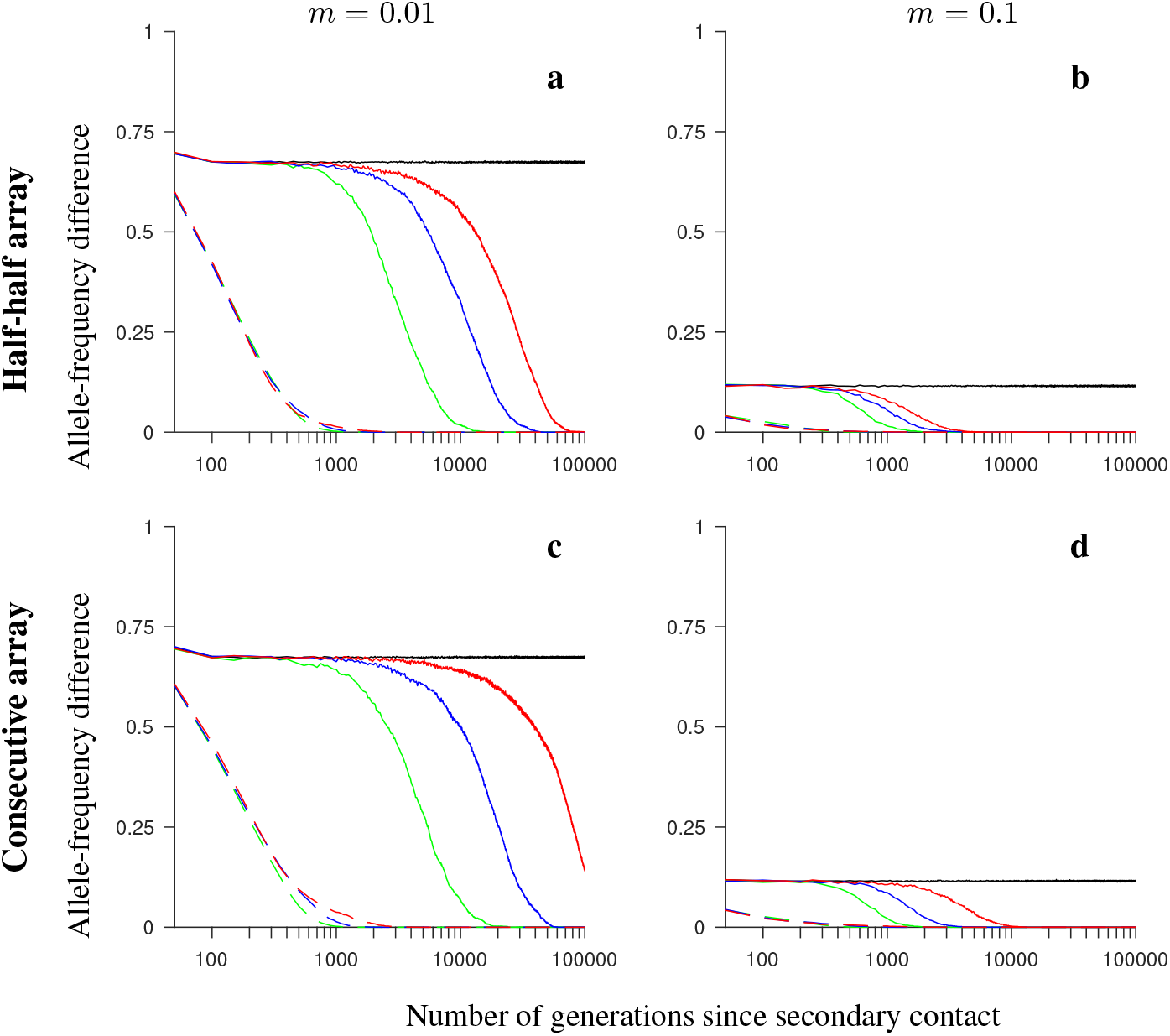
Simulation results for the *conserved-size* model involving locally favored and neutral alleles. The figure shows the allele-frequency difference between the two populations averaged over 200 independent realizations of the model, and over all loci within the region, as a function of the number of generations since secondary contact. Note the logarithmic scale on the horizontal axis. Gene flux in heterokaryotypes is set to *r*_Inv_ = 2 · 10^−4^ (solid colored lines) or to *r*_Inv_ = 0 (black). Dashed lines show the results for the model without inversions. Number of loci within the region: 4 (green), 8 (blue), 20 (red). The panels differ by the migration rate (*m*), and by the ordering of the loci ***A***_*i*_ and ***B***_*i*_, as indicated in the figure. Remaining parameter values: selection strength *s* = 0.1, number of individuals in each population *N* = 1,000, recombination rate in homokaryotypes *r* = 0.1.

**Figure S2.**
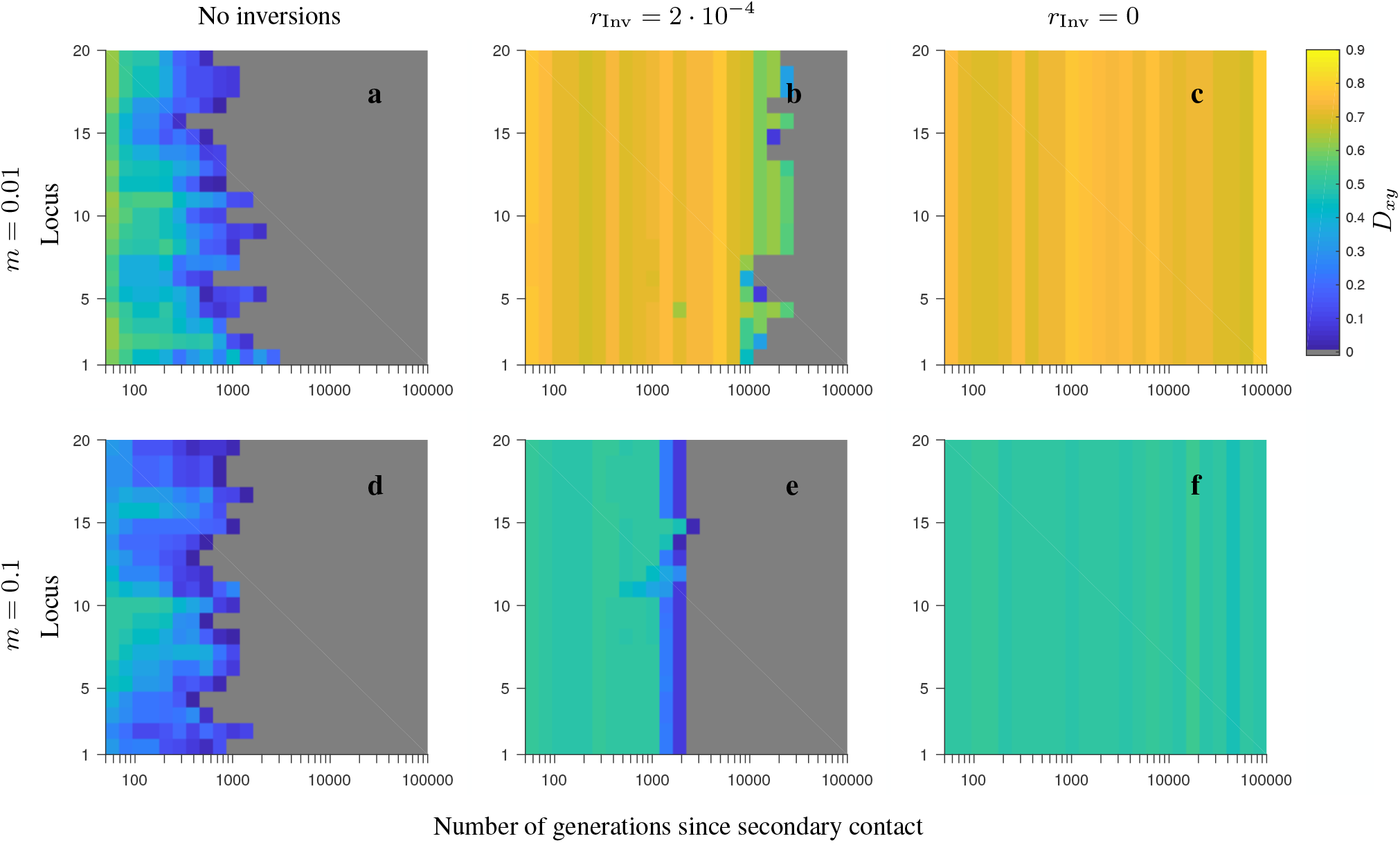
Simulation results for single randomly chosen realizations of the *conserved-size, half-half array* model with 20 loci involving locally favored and neutral alleles. The heat maps depict the evolution of *D*_*xy*_ (see the color map) as a function of the number of generations since secondary contact (horizontal axis), and as a function of the locus number within the region considered (vertical axis). Grey depicts either very low value of *D*_*xy*_, or fixation. Note the logarithmic scale on the horizontal axis. The panels differ by the migration rate (*m*), and by whether or not the model involves inversions, as indicated in the figure. Note that for the model involving inversions, the results are shown for the gene flux in heterokaryotypes set to *r*_Inv_ = 2 · 10^−4^ (second column) or to *r*_Inv_ = 0 (third column). Remaining parameters: number of loci 20, local population size 1,000, selection strength *s* = 0.1, recombination rate in homokaryotypes *r* = 0.1.

**Figure S3.**
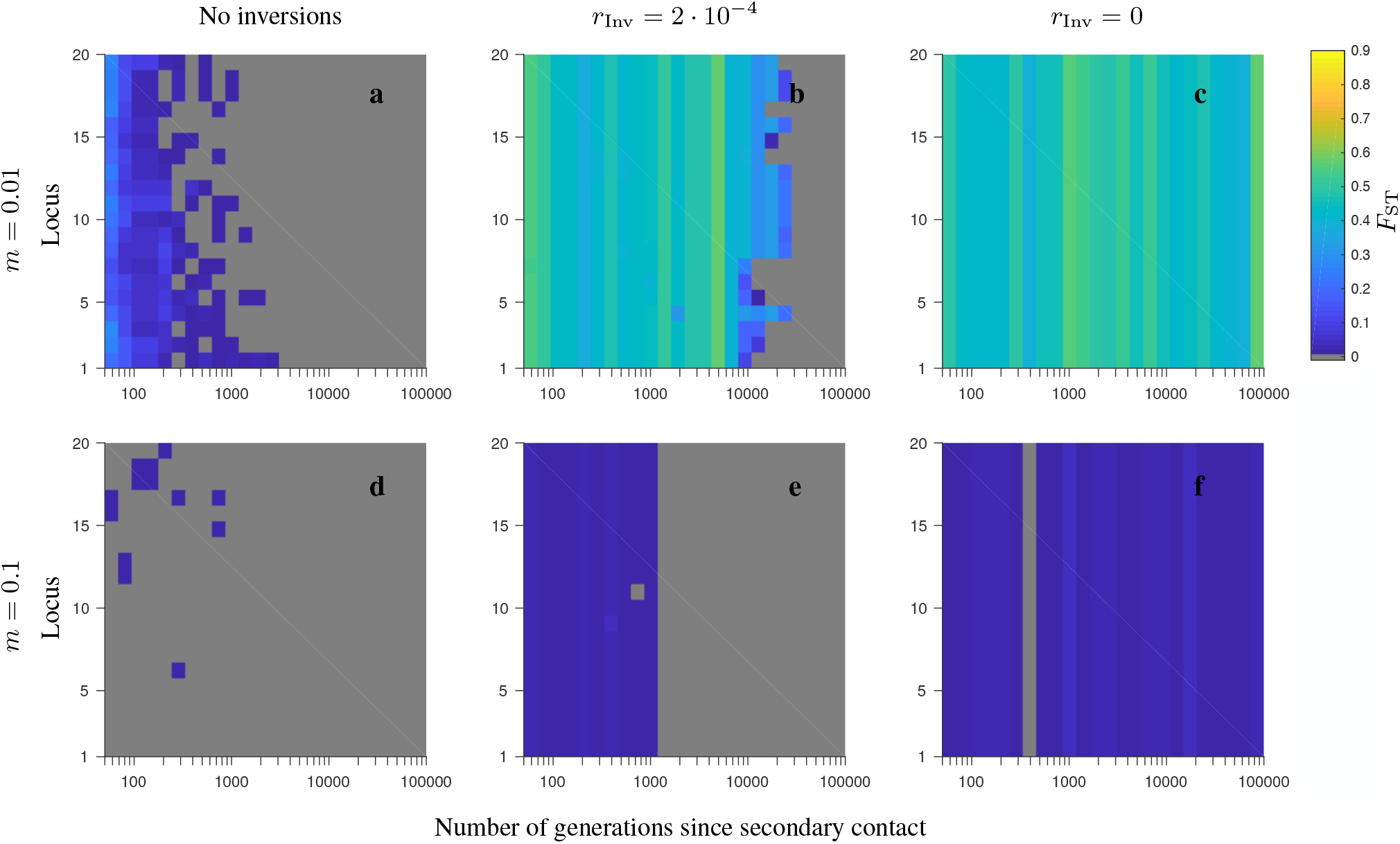
Same as in Fig. S2, but for the corresponding *F*_ST_ patterns.

**Figure S4.**
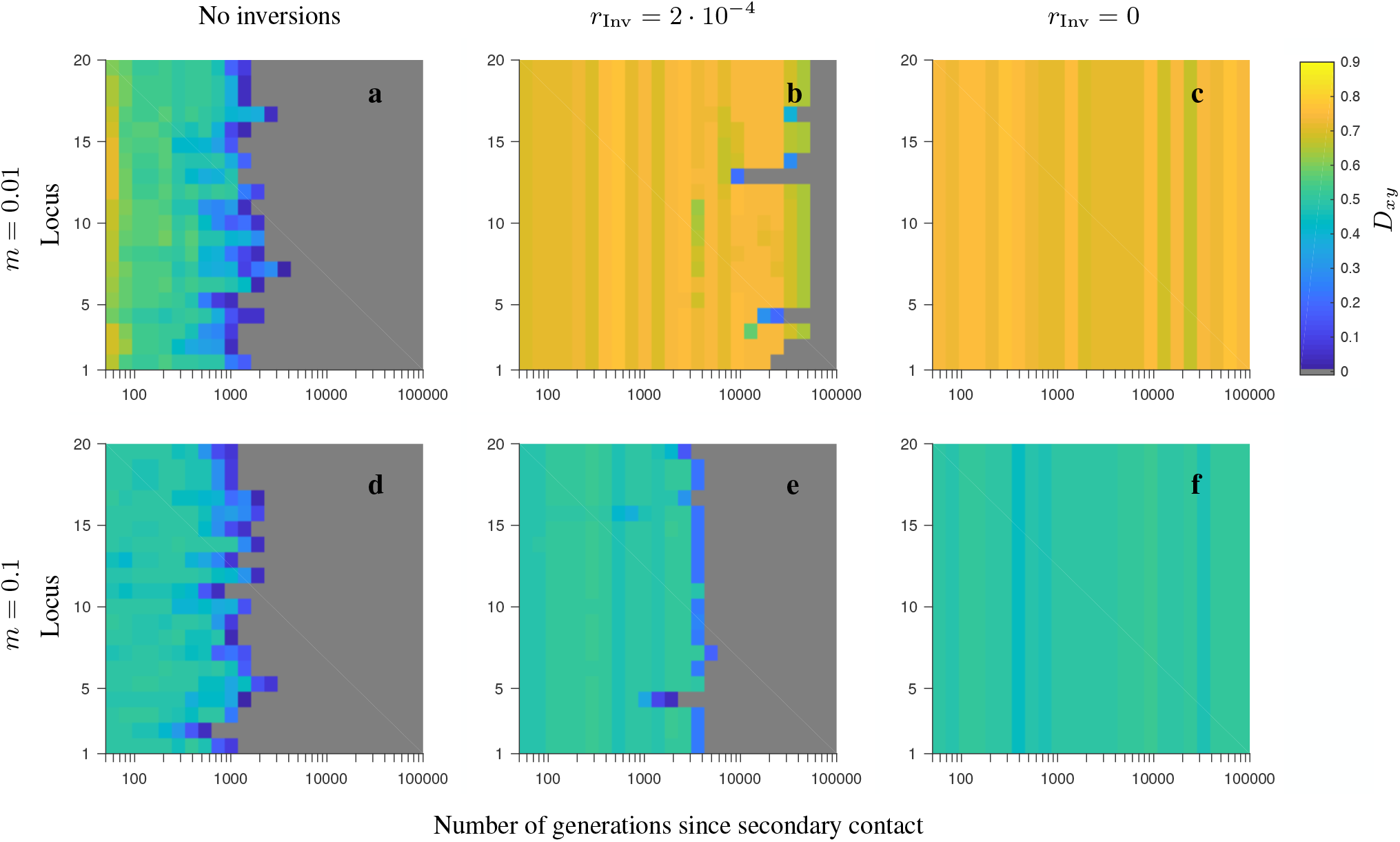
Same as in Fig. S2, but for the *consecutive-array*, *conserved-size* model involving locally favored and neutral alleles.

**Figure S5.**
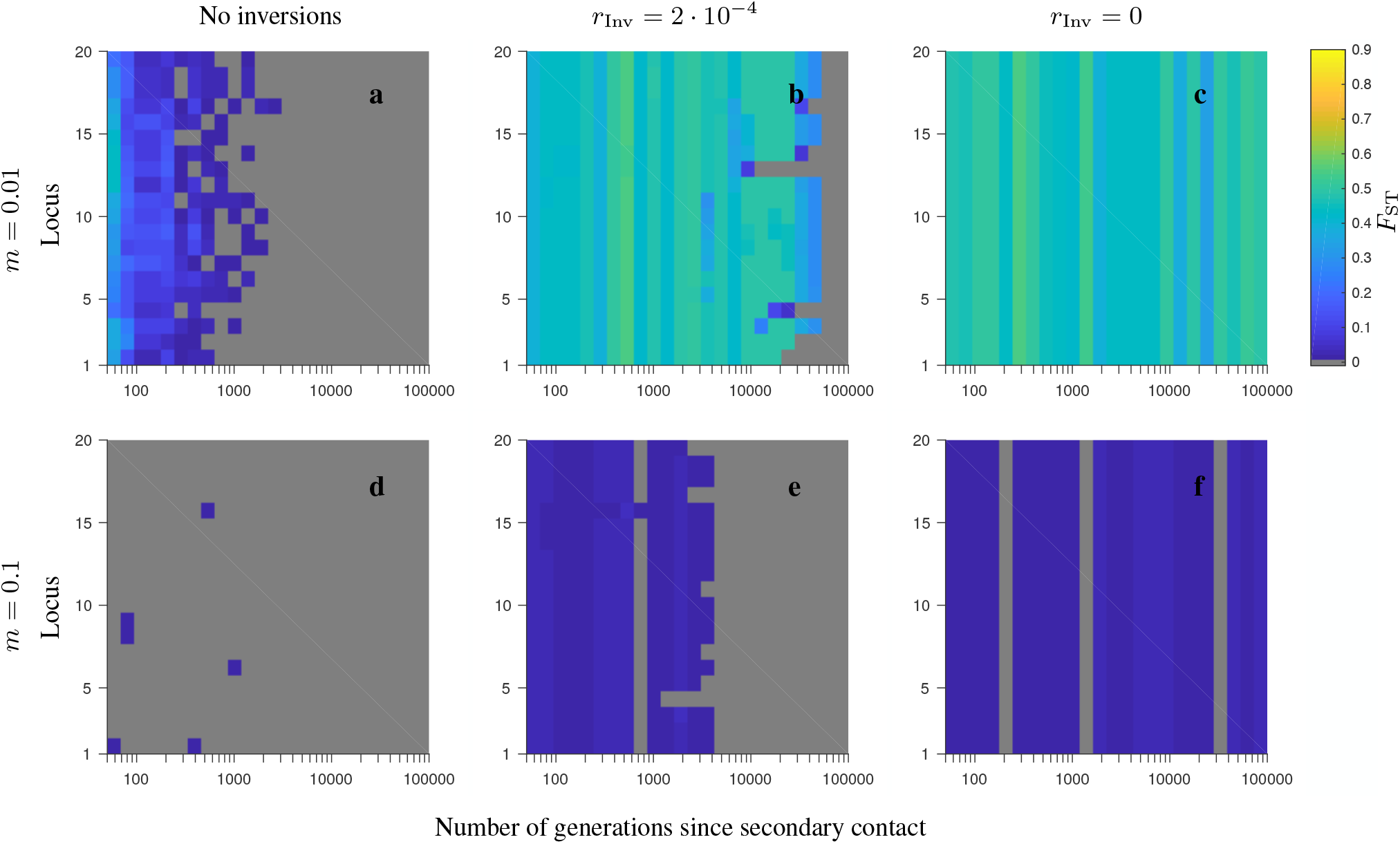
Same as in Fig. S4, but for the corresponding *F*_ST_ patterns.

**Figure S6.**
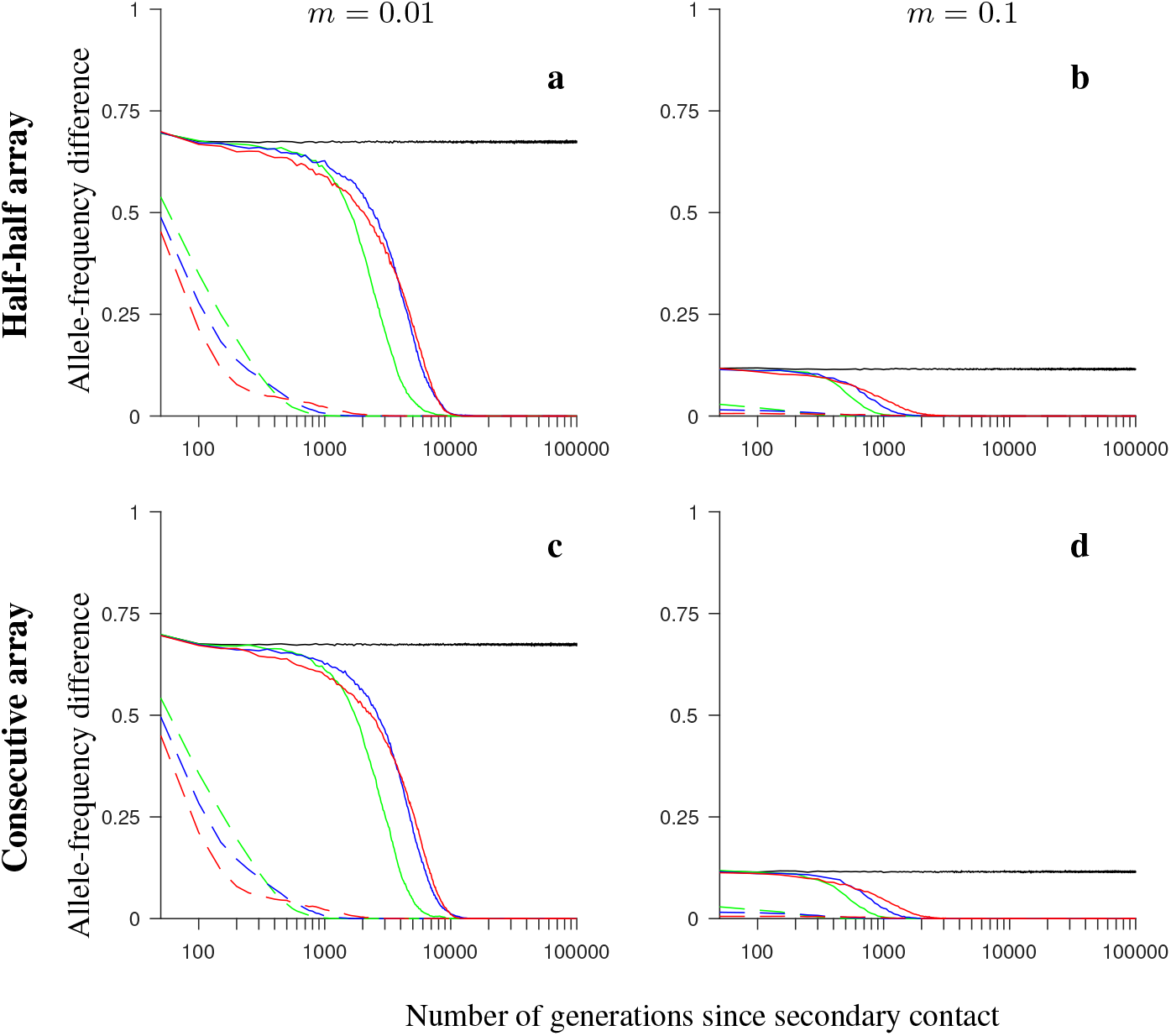
Same as in Fig. S1 but for the *increasing-size* version of the model with locally favored and neutral alleles.

**Figure S7.**
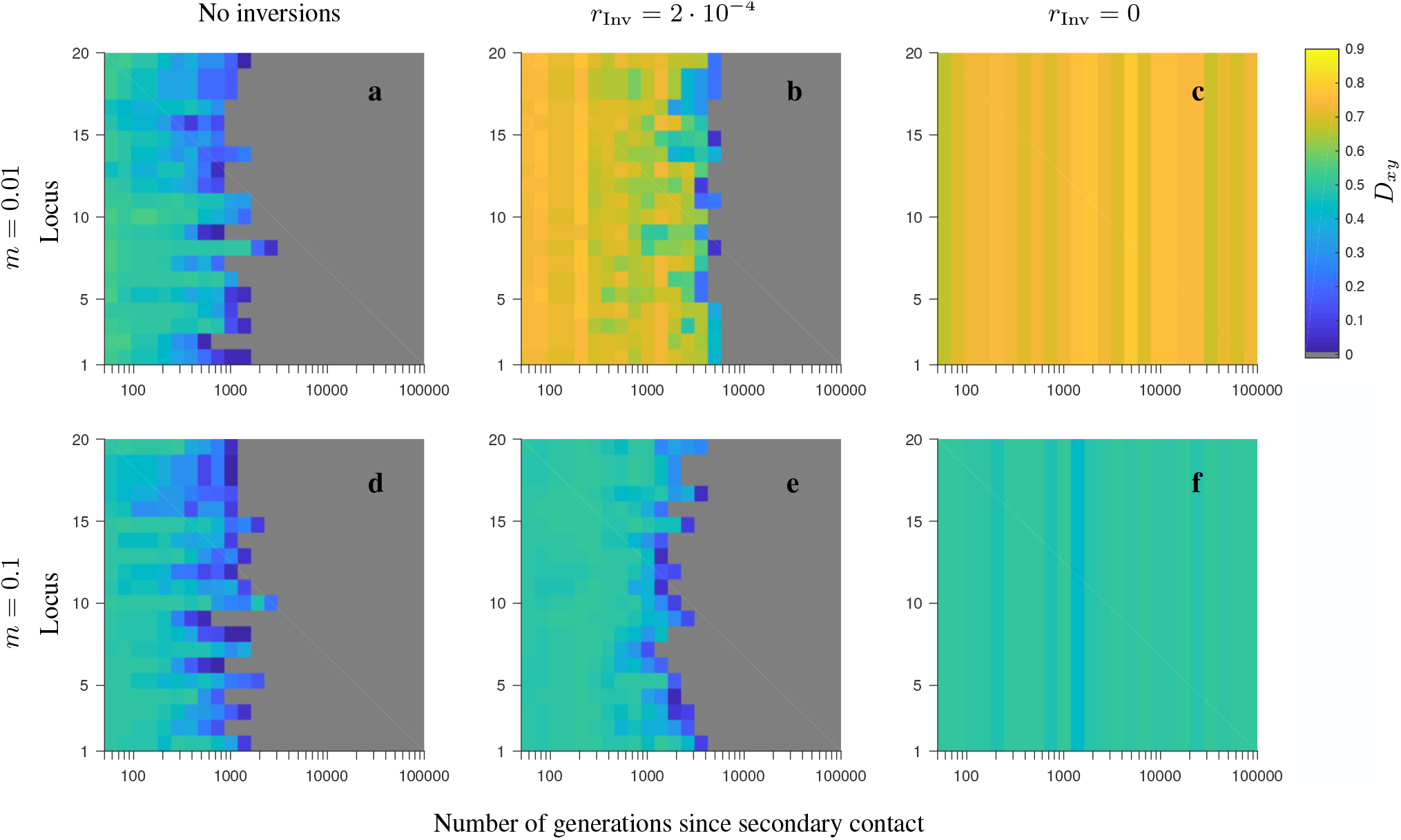
Same as in Fig. S2 but for the *increasing-size* version of the *half-half array* model with locally favored and neutral alleles.

**Figure S8.**
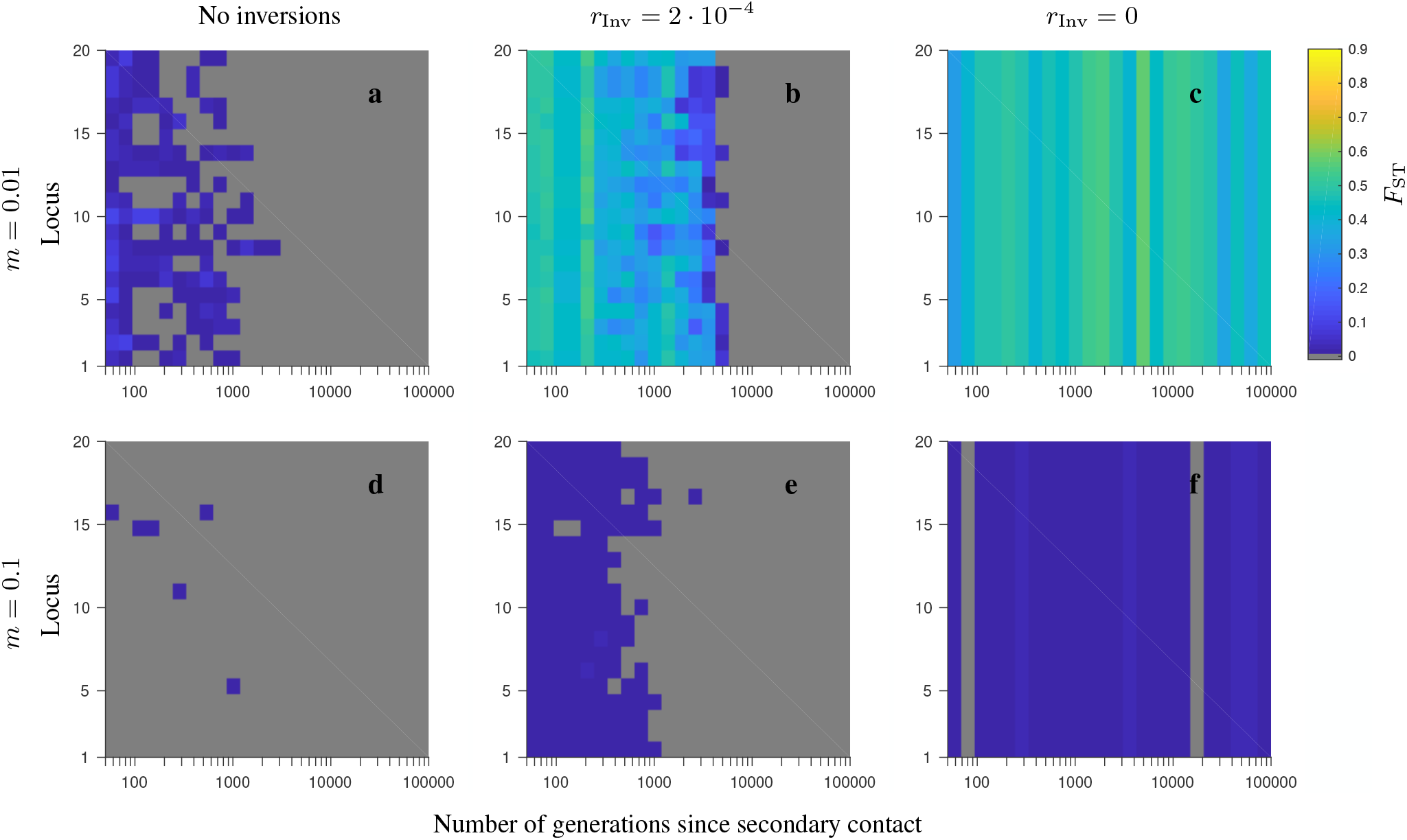
Same as in Fig. S7, but for the corresponding *F*_ST_ patterns.

**Figure S9.**
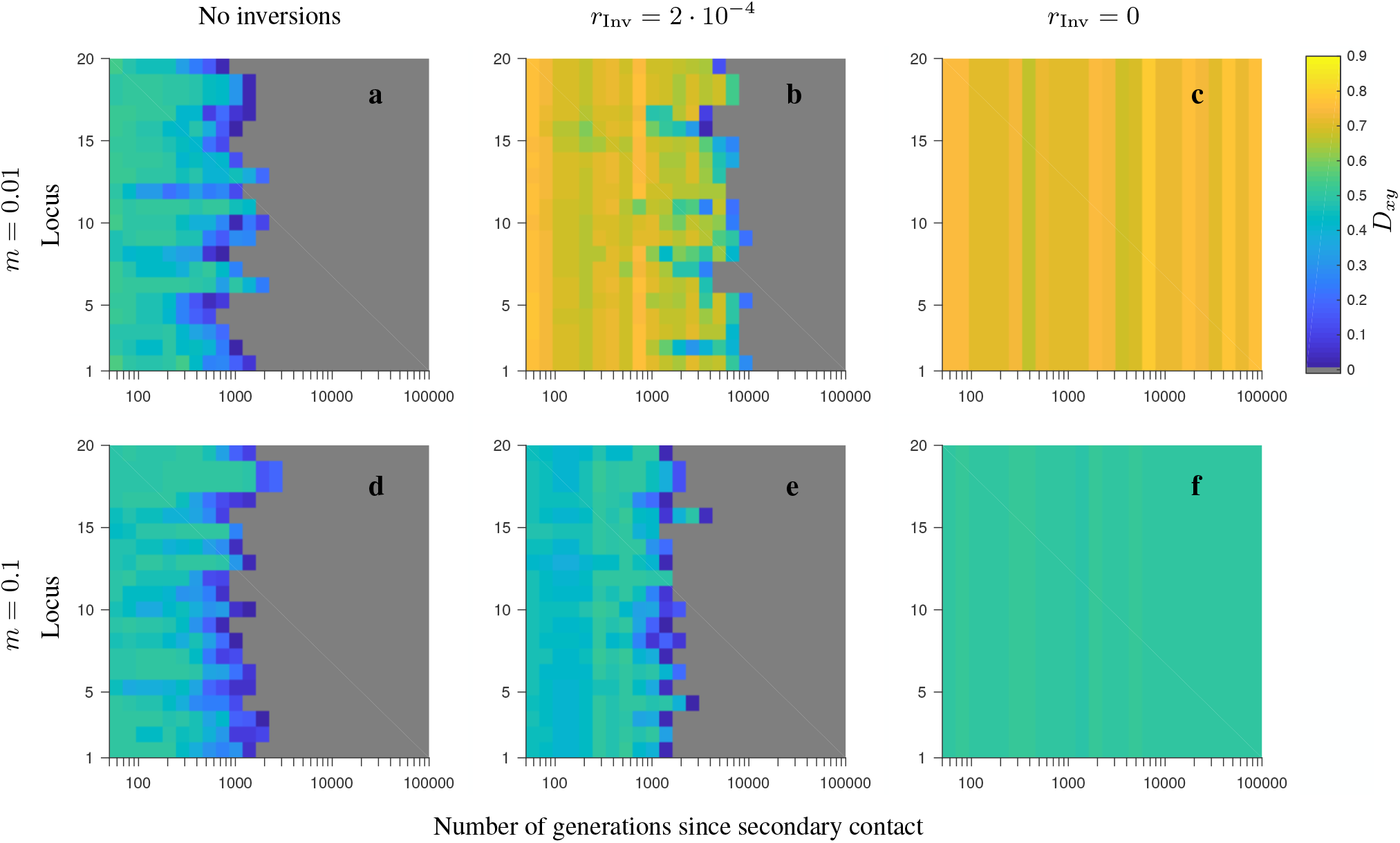
Same as in Fig. S7, but for the *consecutive-array* version of the *increasing-size* model with locally favored and neutral alleles.

**Figure S10.**
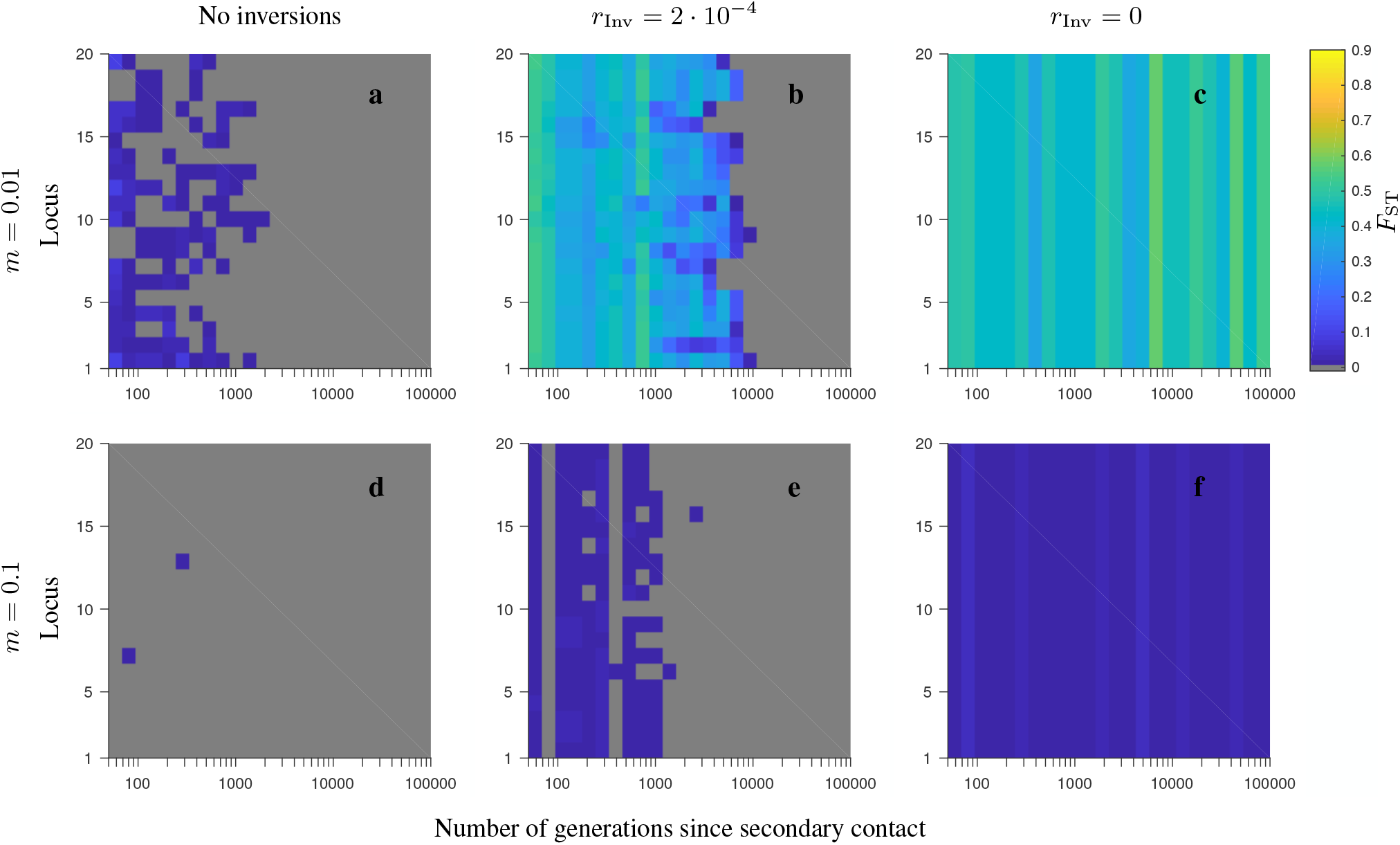
Same as in Fig. S9, but for the corresponding *F*_ST_ patterns.

**Figure S11.**
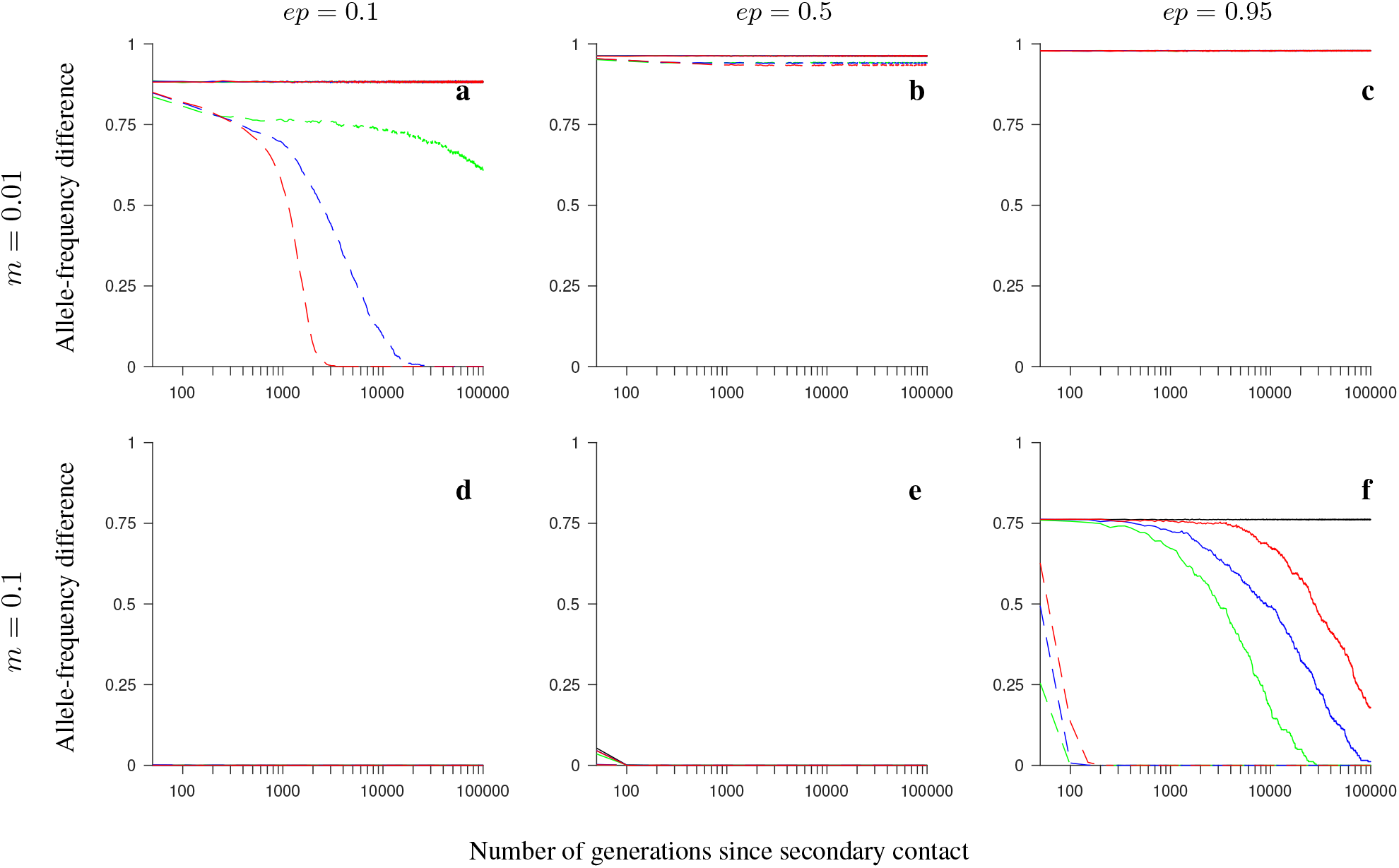
Simulation results for the *half-half array*, *conserved-size* model involving universally beneficial alleles with genetic incompatibilities. The figure shows the allele-frequency difference between the two populations averaged over 200 independent realizations of the model, and over all loci within the region, as a function of the number of generations since secondary contact. Note the logarithmic scale on the horizontal axis. Gene flux in heterokaryotypes is set to *r*_*Inv*_ = 2 · 10^−4^ (colored solid lines) or to *r*_*Inv*_ = 0 (black). Dashed lines show the results for the model without inversions. Number of loci within the region: 4 (green), 8 (blue), 20 (red). Note that all lines overlap in panels **c**-**e**; in addition, solid lines, as well as dashed lines, mutually overlap in panels **a** and **b**. Remaining parameter values: selection strength *s* = 0.1, number of individuals in each population *N* = 1,000, recombination rate in homokaryotypes *r* = 0.1.

**Figure S12.**
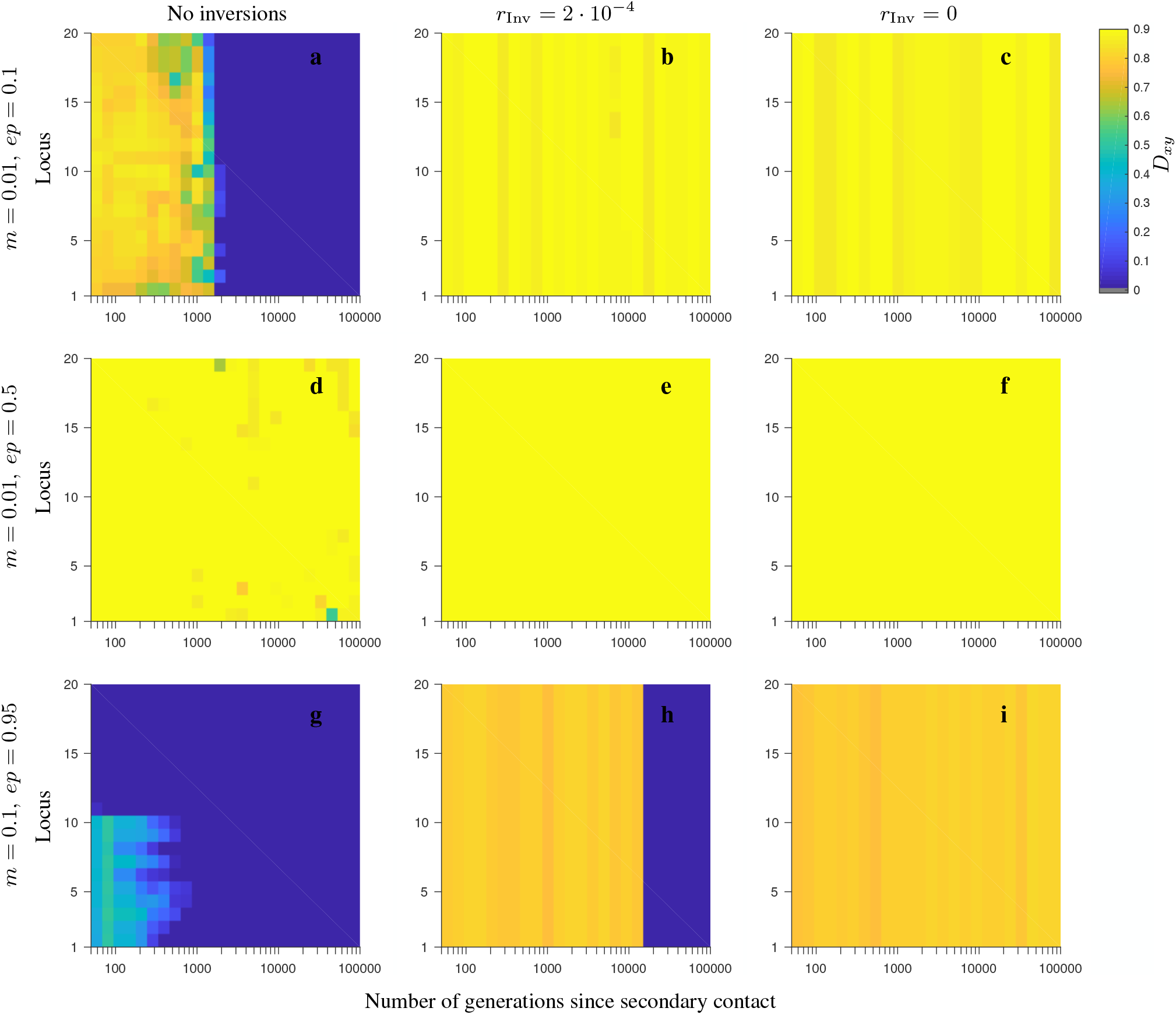
Simulation results for single randomly chosen realizations of the *conserved-size, half-half array* model with 20 loci involving universally beneficial alleles with genetic incompatibilities. The heat maps depict the evolution of *D*_*xy*_ (see the color map) as a function of the number of generations since secondary contact (horizontal axis), and as a function of the locus number within the region considered (vertical axis). Note the logarithmic scale on the horizontal axis. The panels differ by the migration rate (*m*), and by the strength of negative epistasis (*ep*), as indicated in the figure. Note that for the model involving inversions, the results are shown for the gene flux in heterokaryotypes set to *r*_Inv_ = 2 · 10^−4^ (second column) or to *r*_Inv_ = 0 (third column). Remaining parameters: number of loci 20, local population size 1,000, selection strength *s* = 0.1, recombination rate in homokaryotypes *r* = 0.1.

**Figure S13.**
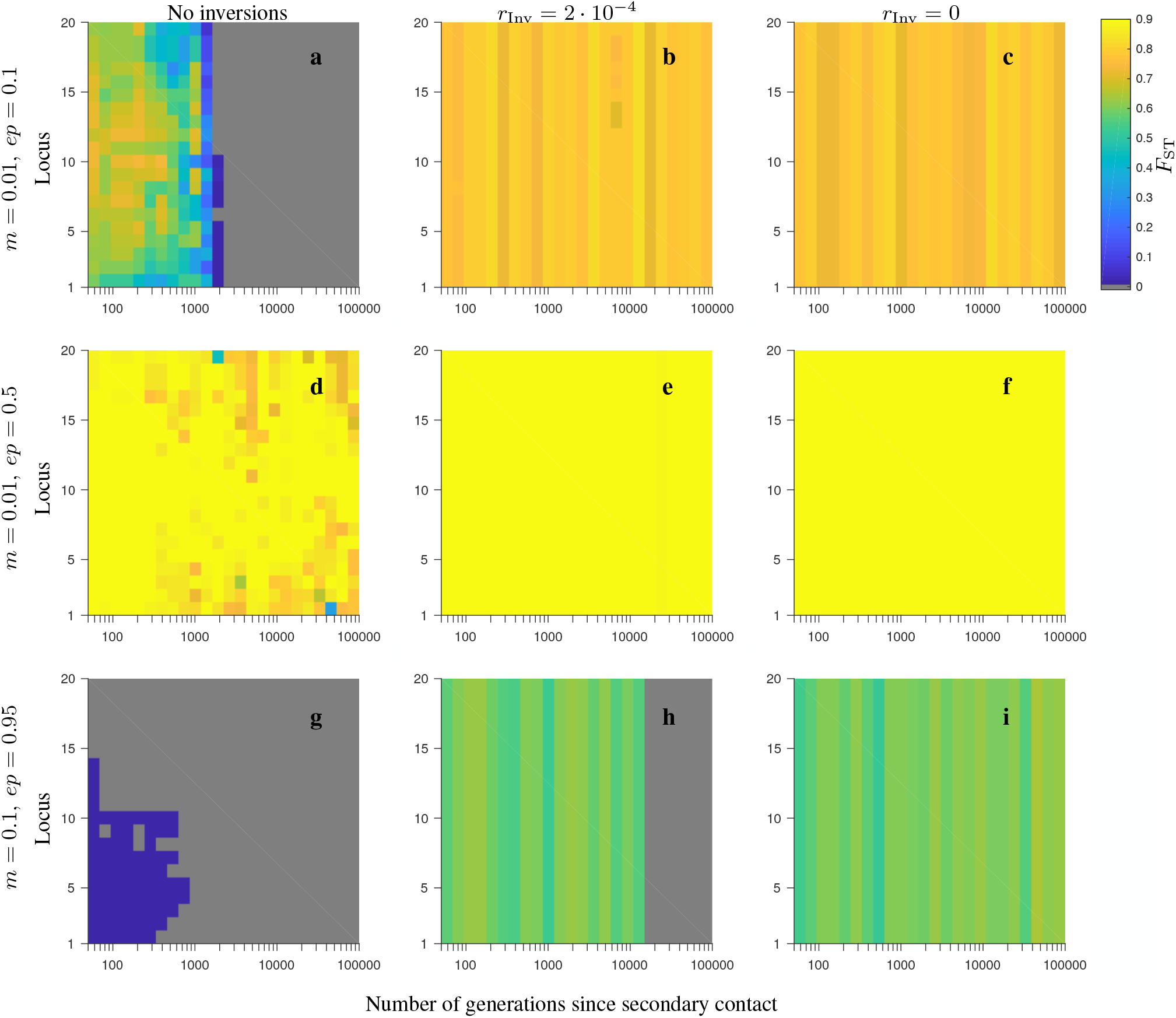
Same as in Fig. S12, but for the corresponding *F*_ST_ patterns.

**Figure S14.**
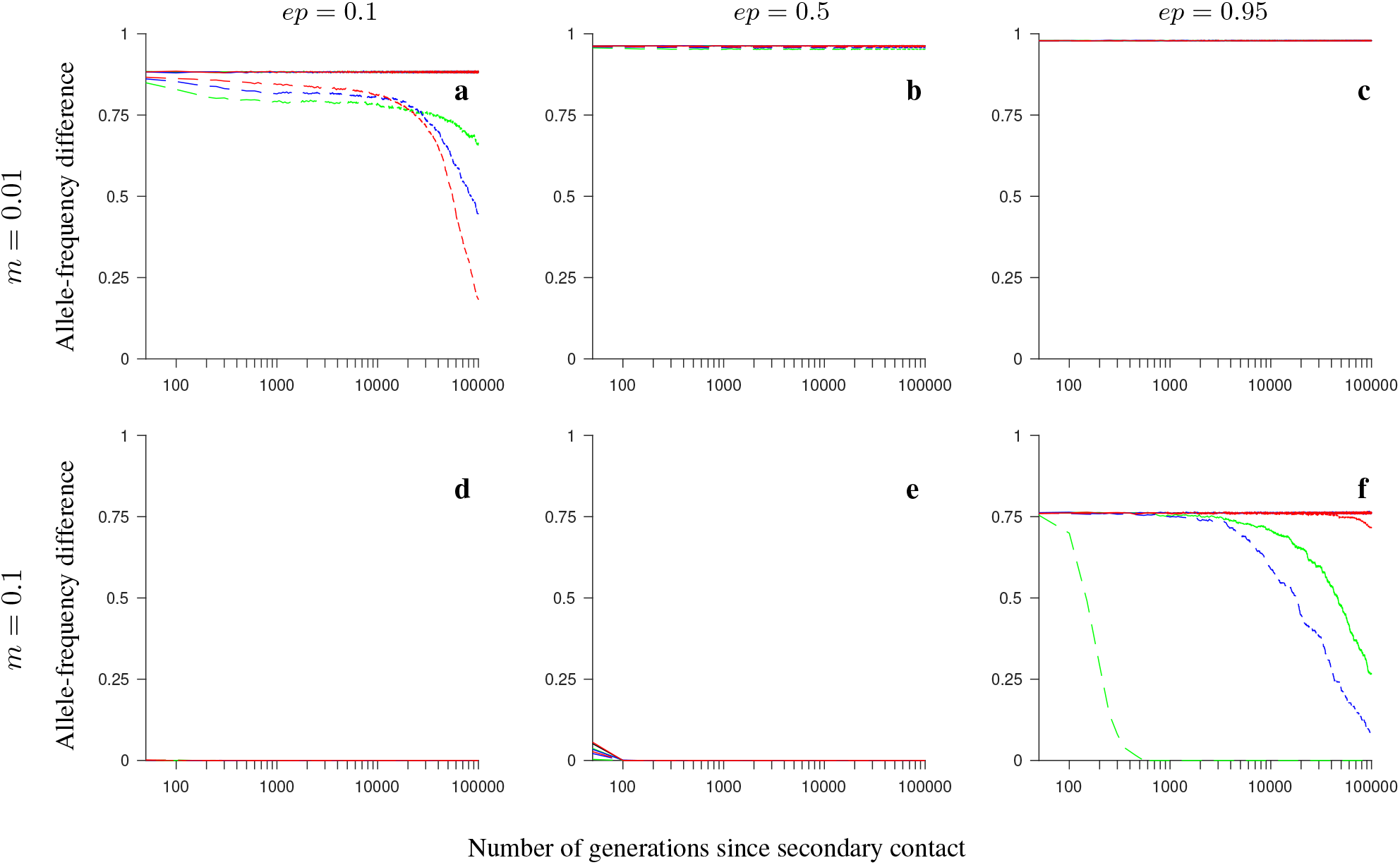
Same as in Fig. S11, but for the *consecutive-array* version of the *conserved-size* model involving universally beneficial alleles with genetic incompatibilities.

**Figure S15.**
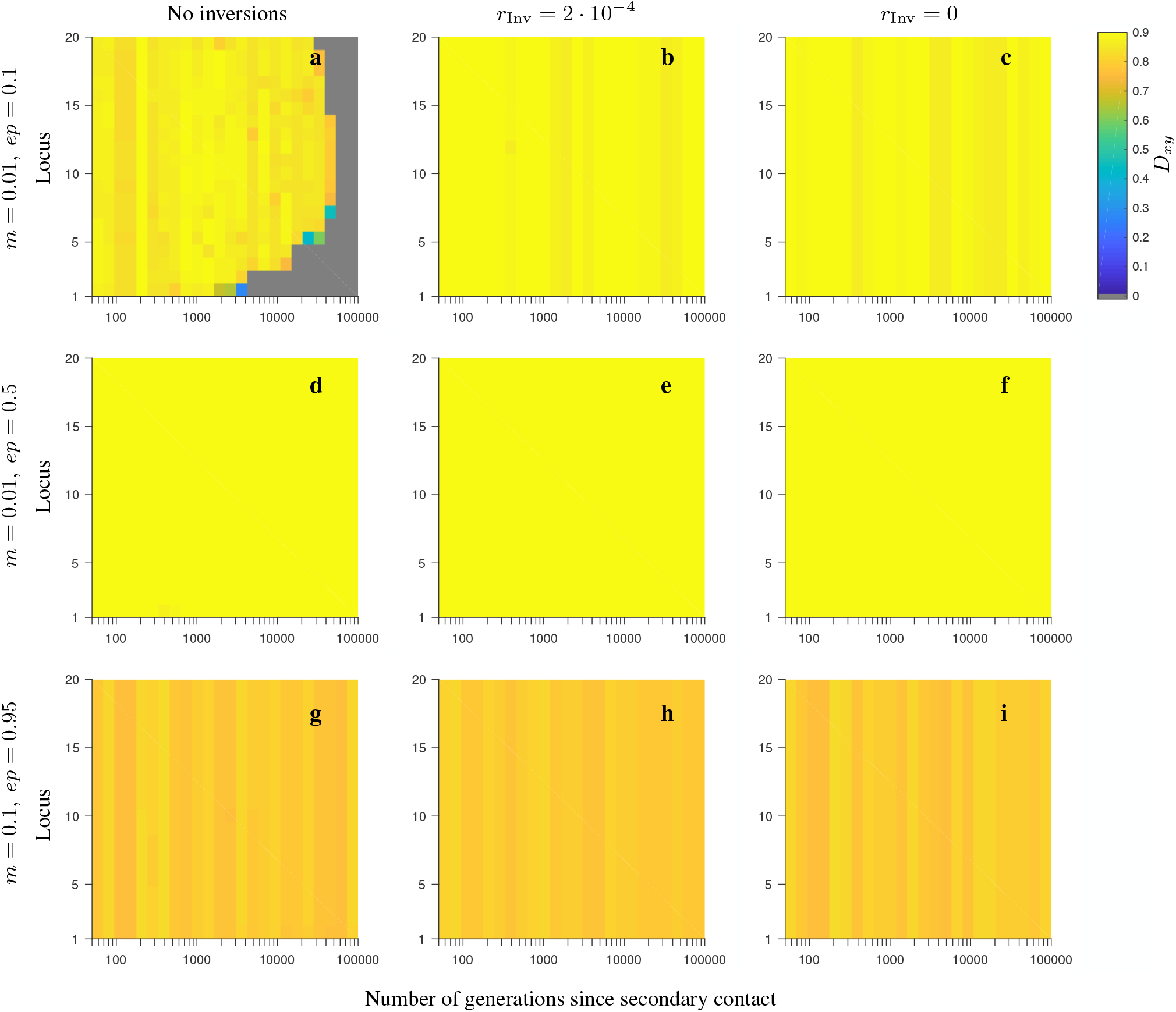
Same as in Fig. S12, but for the *consecutive-array* version of the *conserved-size* model involving universally beneficial alleles with genetic incompatibilities.

**Figure S16.**
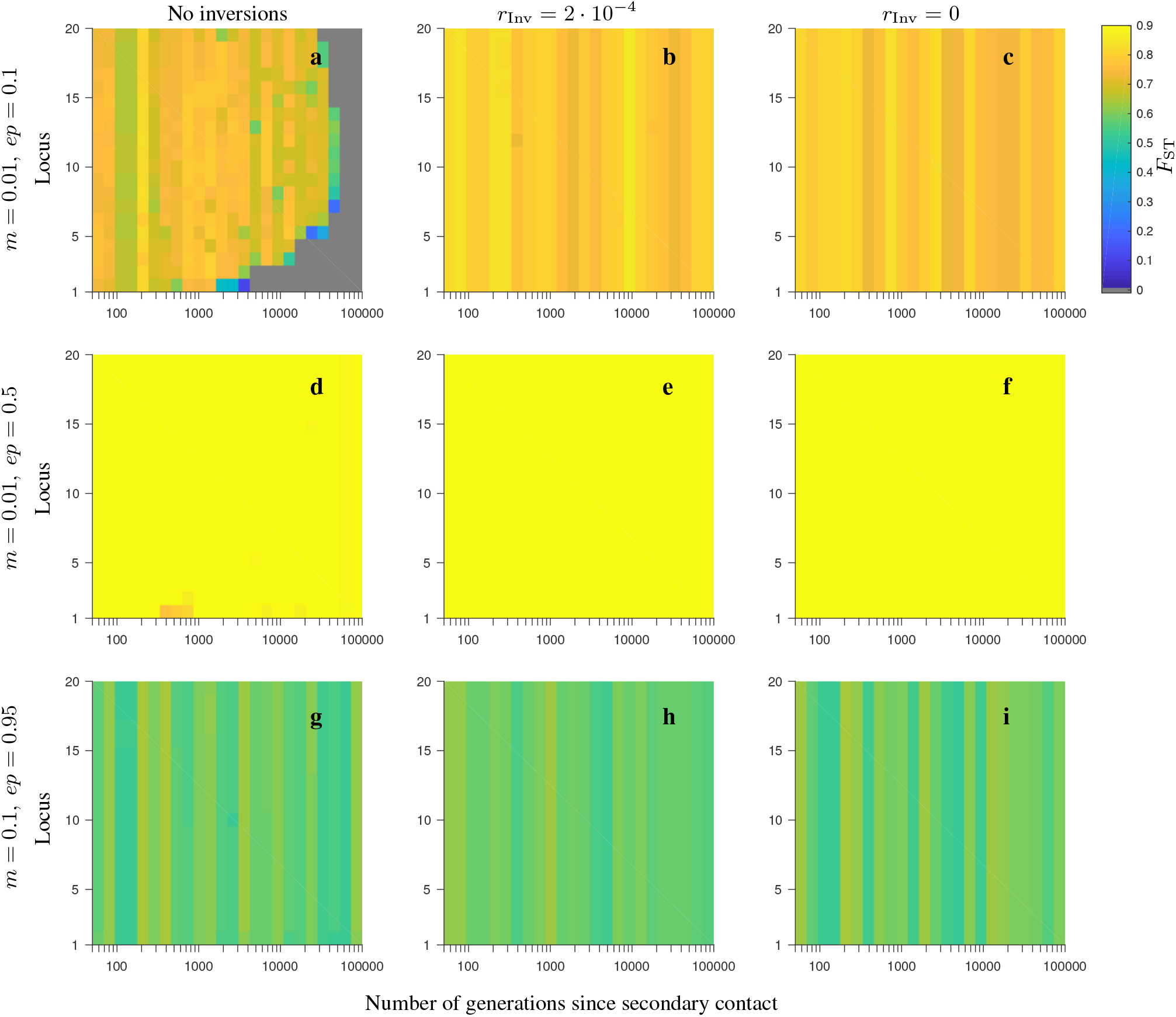
Same as in Fig. S15, but for the corresponding *F*_ST_ patterns.

**Figure S17.**
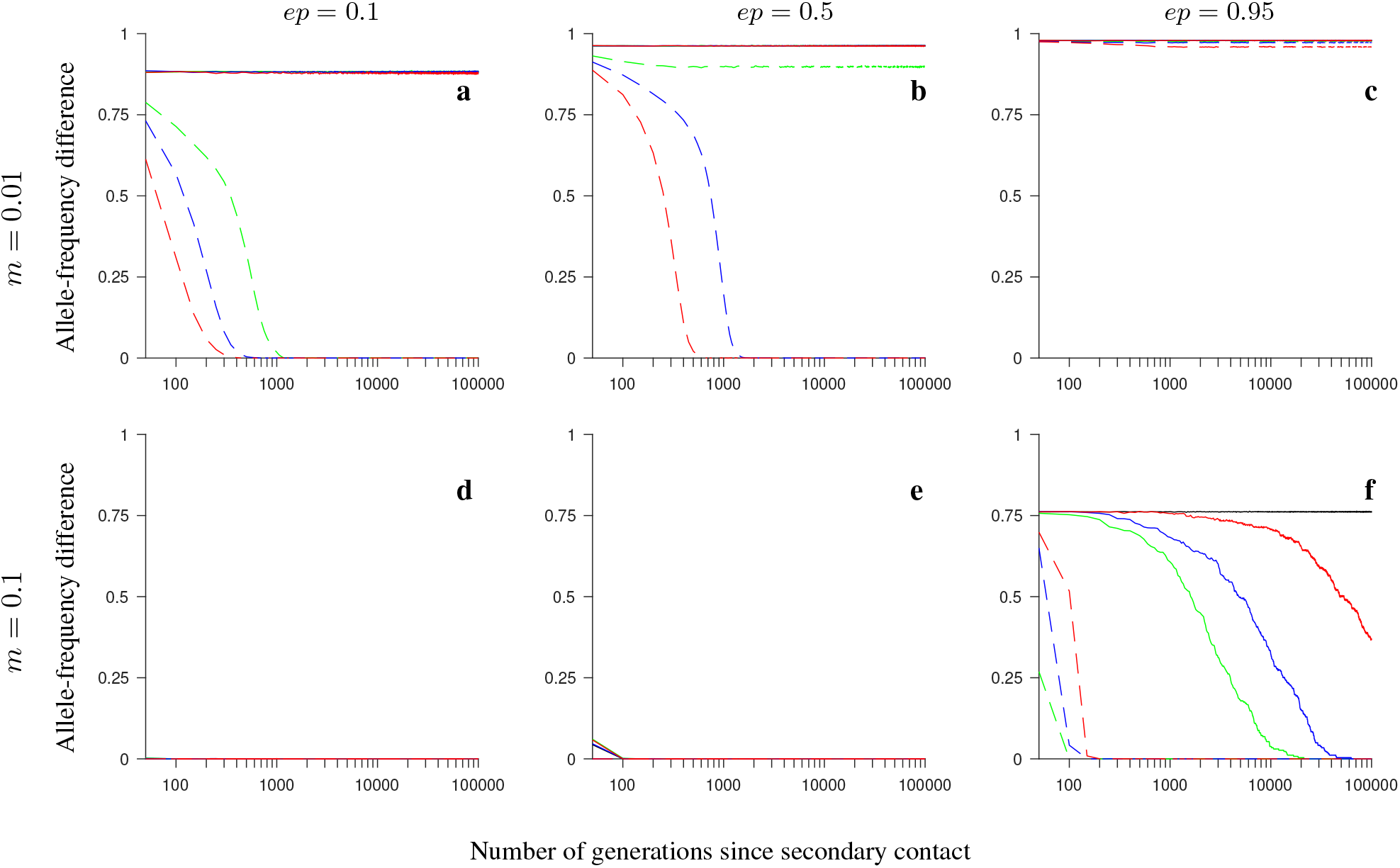
Same as in Fig. S11, but for the *increasing-size* version of the *half-half array* model involving universally beneficial alleles with genetic incompatibilities.

**Figure S18.**
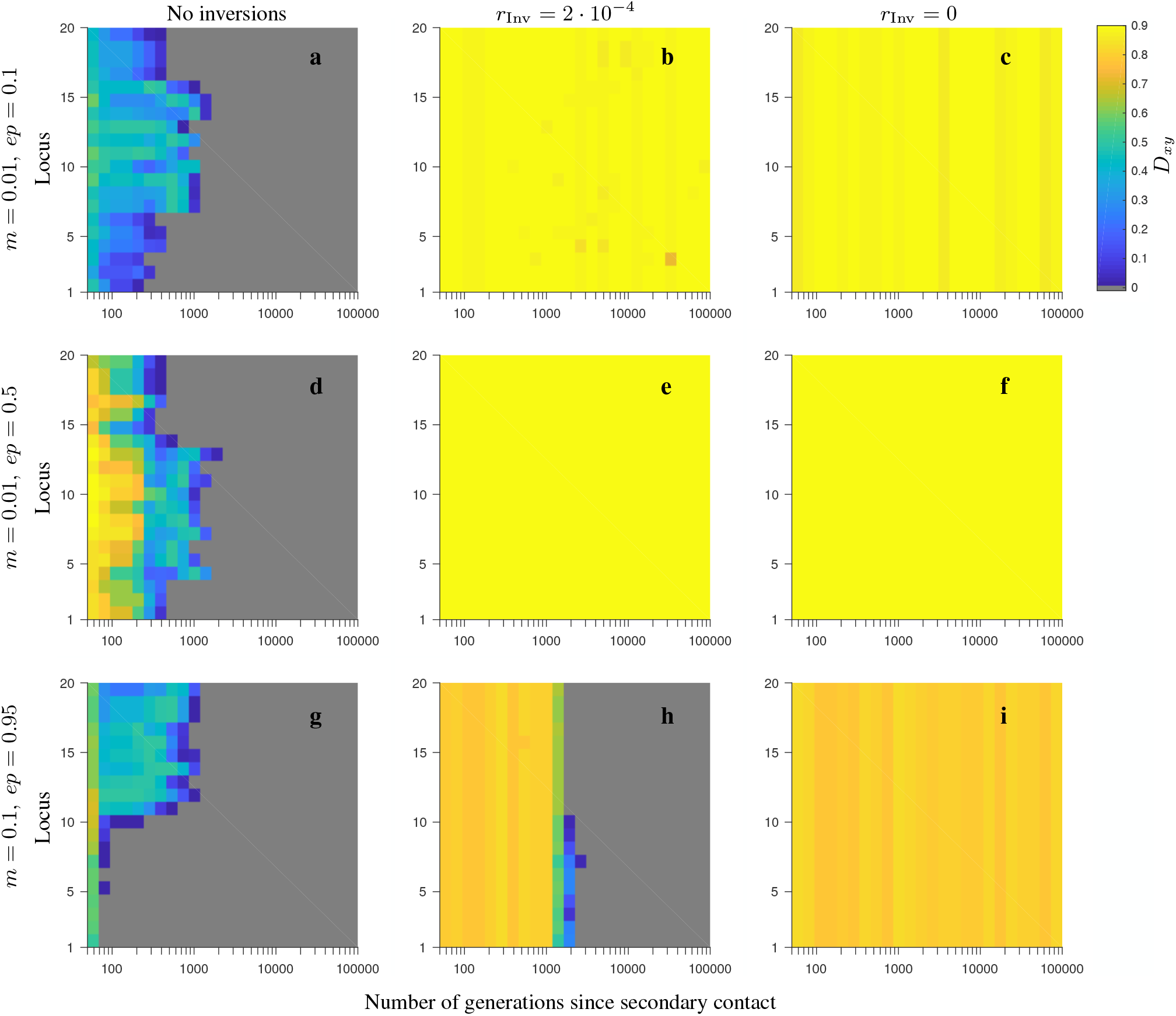
Same as in Fig. S12, but for the *increasing-size* version of the *half-half array* model involving universally beneficial alleles with genetic incompatibilities.

**Figure S19.**
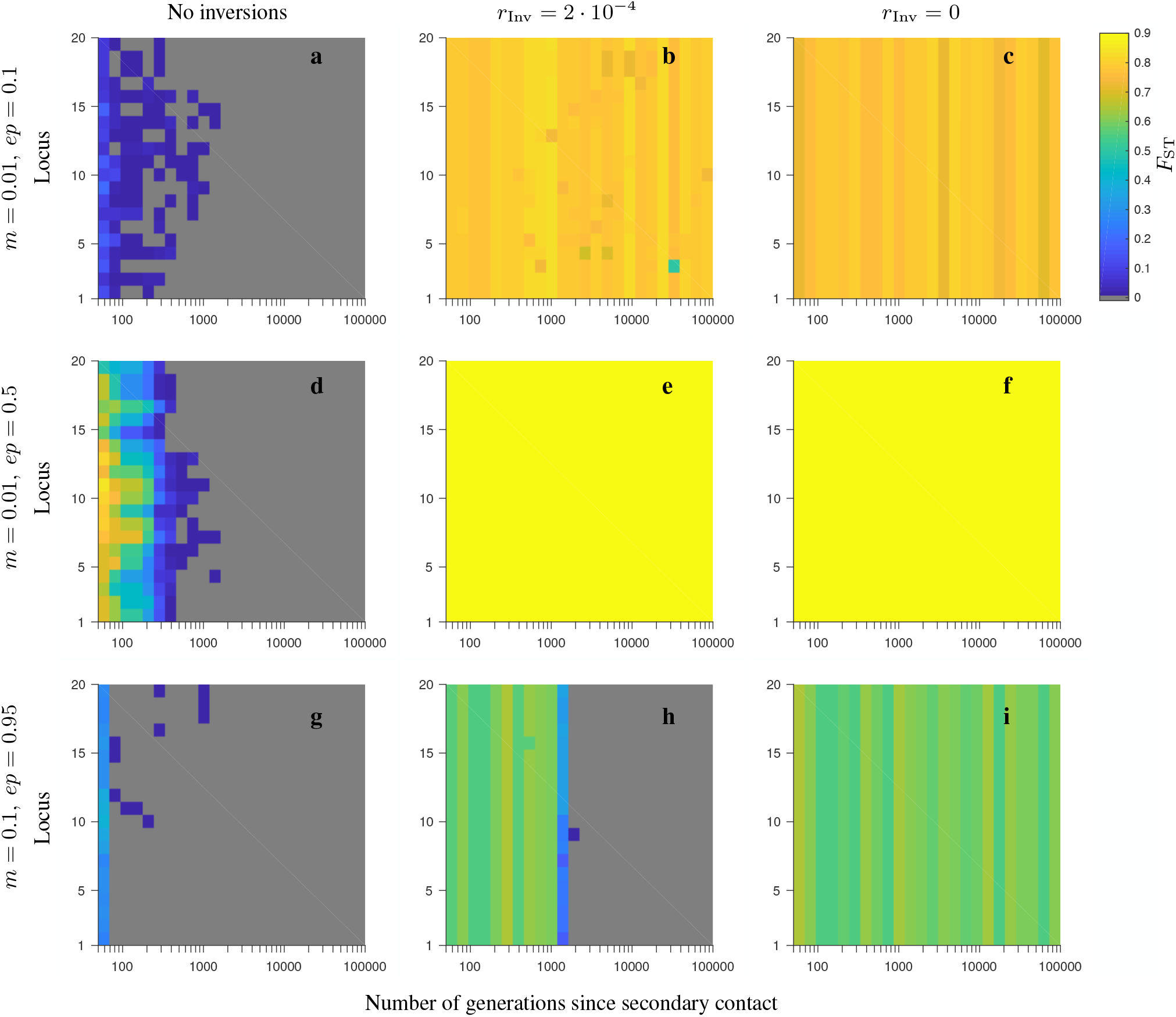
Same as in Fig. S18, but for the corresponding *F*_ST_ patterns.

**Figure S20.**
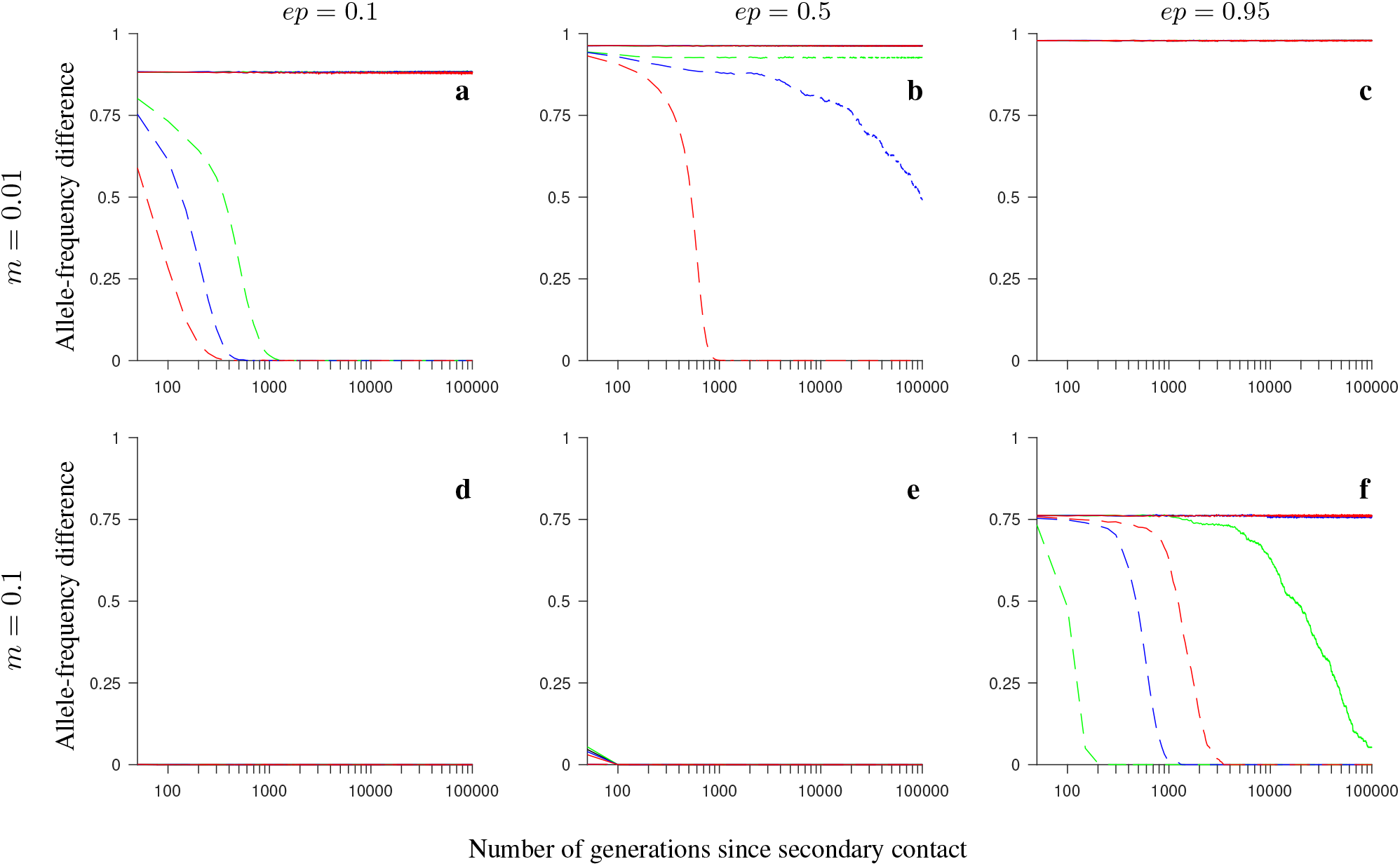
Same as in Fig. S11, but for the *increasing-size* version of the *consecutive-array* model involving universally beneficial alleles with genetic incompatibilities.

**Figure S21.**
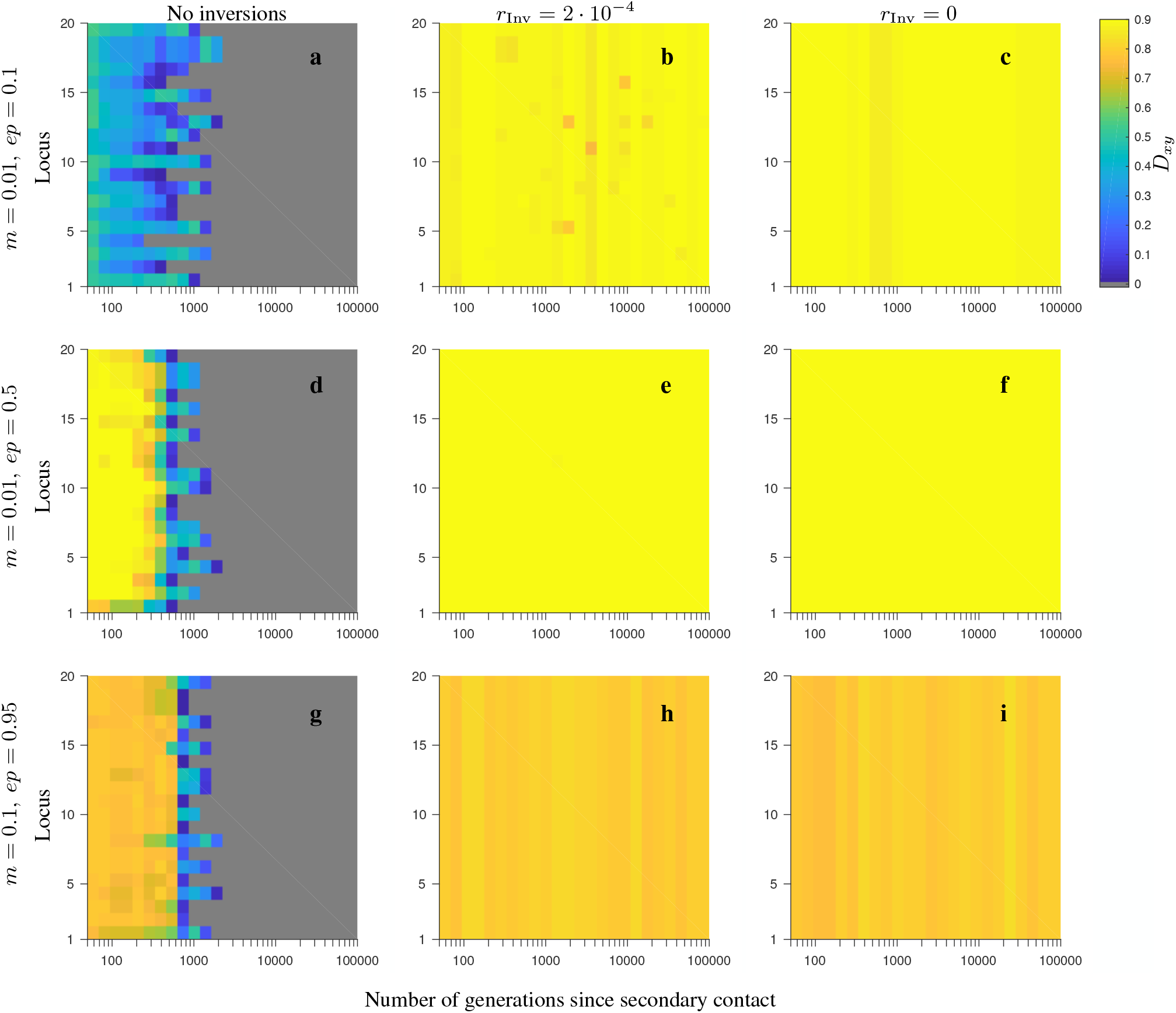
Same as in Fig. S12, but for the *increasing-size* version of the *consecutive-array* model involving universally beneficial alleles with genetic incompatibilities.

**Figure S22.**
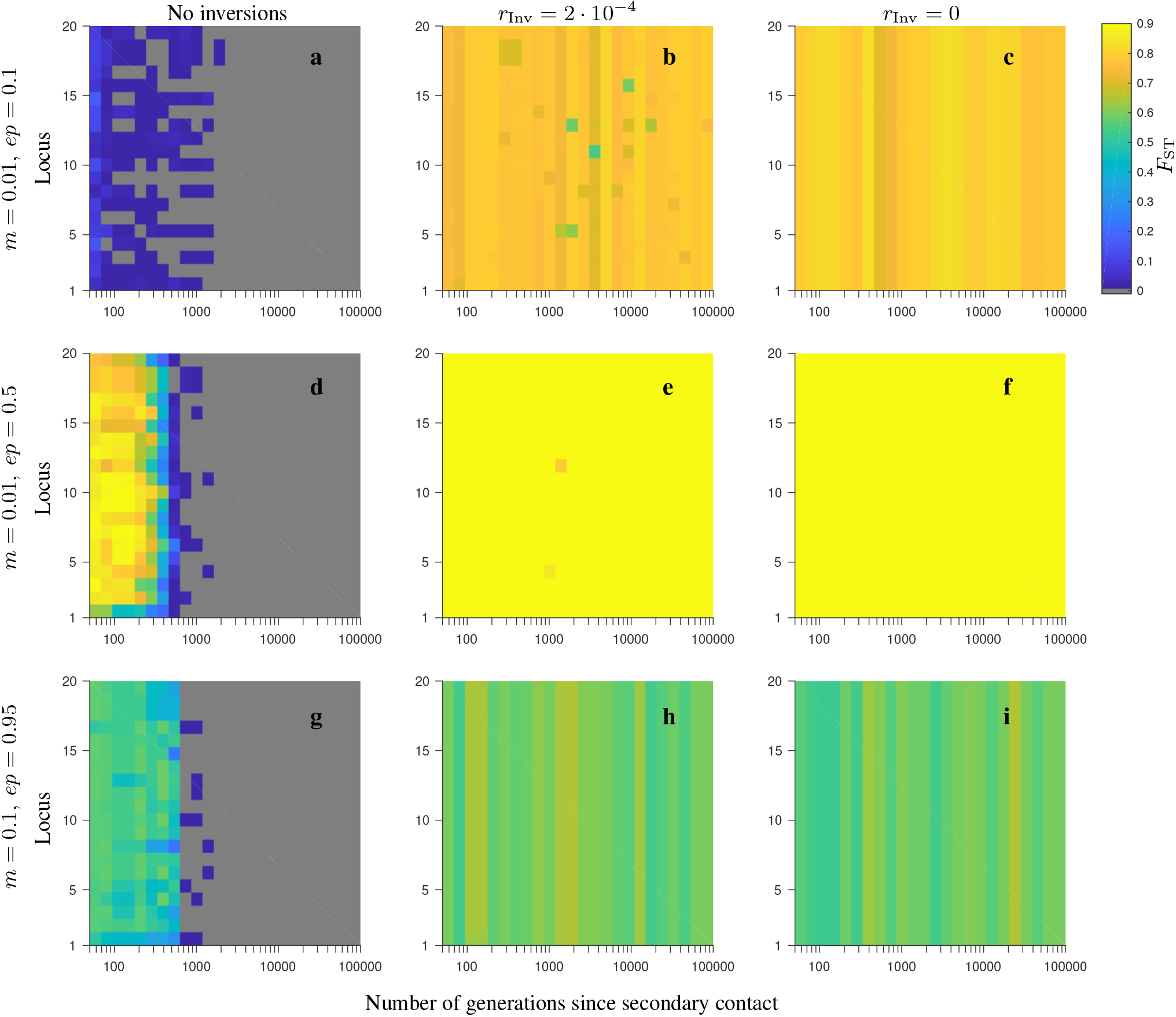
Same as in Fig. S21, but for the corresponding *F*_ST_ patterns.

**Figure S23.**
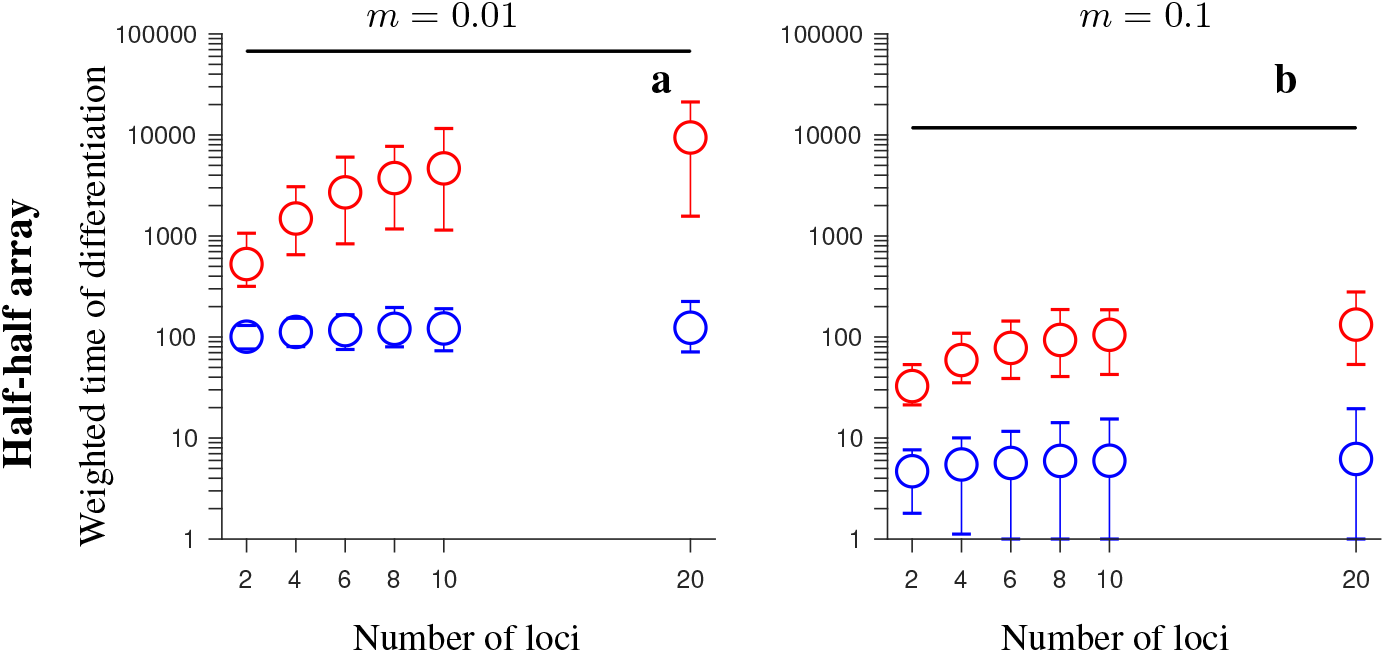
The effect of population size: simulation results for the *conserved-size half-half array* model involving locally favored and neutral alleles, and with local population size set to *N* = 5,000. The figure shows the weighted time of differentiation (*T*_*w*_) averaged over 200 independent realizations of the model, and over all loci within the region, as a function of the number of loci. Note the logarithmic scale on the vertical axis. Gene flux in heterokaryotypes is set to *r*_*Inv*_ = 2 · 10^−4^ (red) or *r*_*Inv*_ = 0 (black). Results for the model without inversions are shown in blue. The vertical lines around the symbols, and the grey regions (for *r*_Inv_ = 0) depict the range between the minimum and maximum values of *T*_*w*_ obtained in individual simulations. The panels differ by the migration rate (*m*) as indicated in the figure. Remaining parameter values: selection strength *s* = 0.1, recombination rate in homokaryotypes *r* = 0.1.

**Figure S24.**
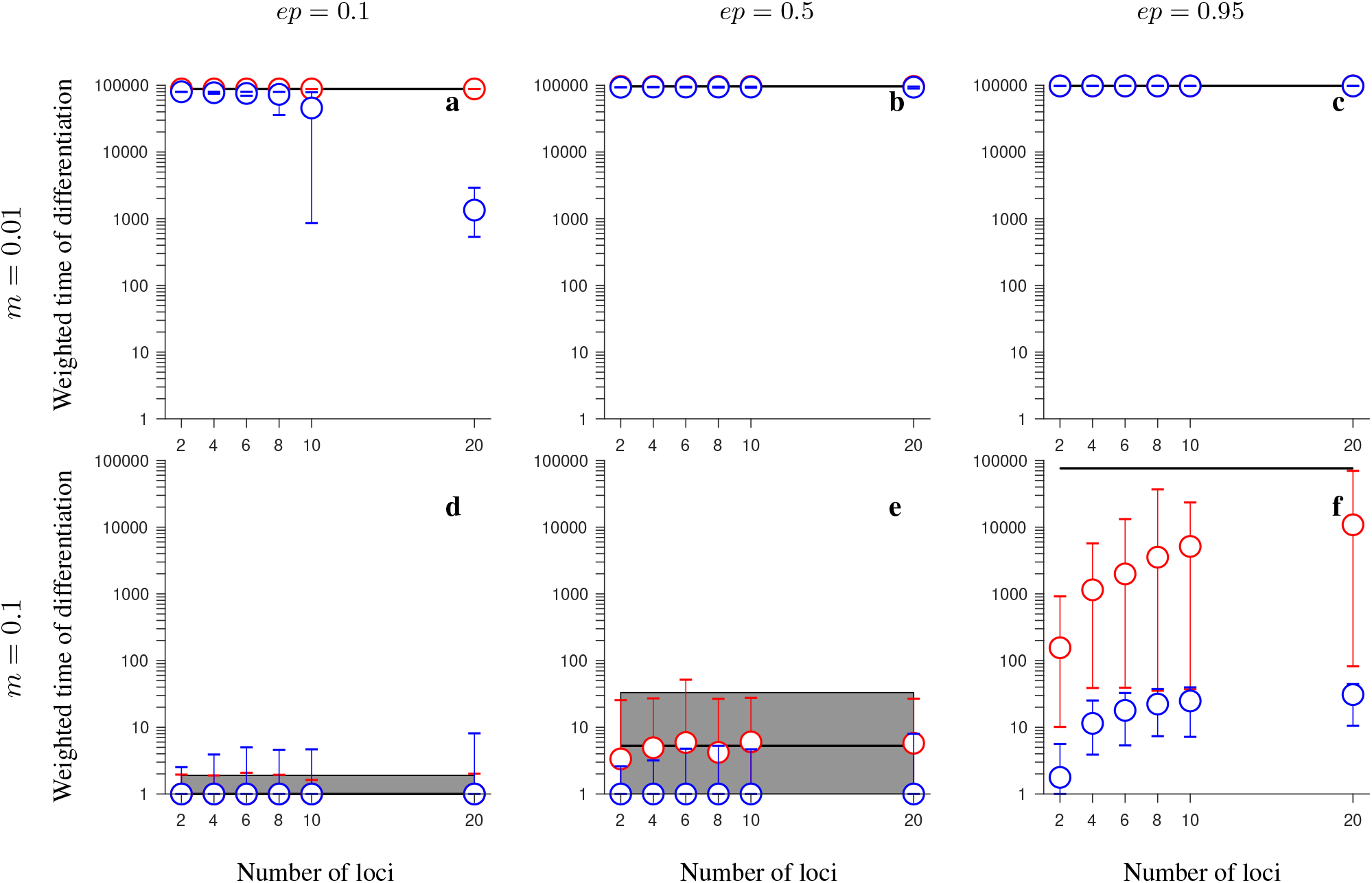
The effect of population size: simulation results for the *conserved-size*, *half-half array* model involving universally beneficial alleles with genetic incompatibilities, and with local population size set to *N* = 5,000. The figure shows the weighted time of differentiation (*T*_*w*_) averaged over 200 independent realizations and over all loci within the region, as a function of the number of loci. Note the logarithmic scale on the vertical axis. Gene flux in heterokaryotypes is set to *r*_Inv_ = 2 · 10^−4^ (red) or *r*_Inv_ = 0 (black). Results for the model without inversions are shown in blue. Note that red and blue circles overlap in panels **b**-**d**. The vertical lines around the symbols, and the grey regions (for *r*_Inv_ = 0) depict the range between the minimum and maximum values of *T*_*w*_ obtained in individual simulations. The panels differ by the migration rate (*m*), and by the strength of negative epistasis (*ep*), as indicated in the figure. Remaining parameter values: selection strength *s* = 0.1, recombination rate in homokaryotypes *r* = 0.1.

